# Dynamic Models of Neural Population Dynamics

**DOI:** 10.1101/2024.10.04.616750

**Authors:** Han Hao, Kai Zhang, Momiao Xiong

## Abstract

The recent developments in artificial intelligence (AI) increase the hope that AI can provide a powerful tool to facilitate scientific discovery and to generate and validate new ideas for scientific research autonomously. Large Language Models (LLMs), such as ChatGPT4 have demonstrated remarkable capabilities in understanding and generating human-like text. Their potential extends beyond simple language tasks, offering transformative possibilities in scientific research of all fields. By leveraging vast amounts of data and advanced computational power, LLMs can assist researchers in generating novel ideas, automating routine tasks, and fostering interdisciplinary collaborations. On September 12, 2024, OpenAI released their updated generative artificial intelligence system called ChatGPTo1. This new AI system, built upon chain-of-thought and reinforcement learning, has greatly enhanced logical reasoning abilities and can effectively solve various complex problems from elementary-level mathematical problems to modern scientific research issues in physics, chemistry, and biology. Unlike previous LLMs in which logical reasoning and data analysis abilities are developed through training on actual data, ChatGPTo1 logical reasoning ability and capacity to generate new scientific ideas are primarily acquired through chain-of-thought processes and reinforcement learning rather than pre-training. To examine this, we specifically tested ChatGPTo1 current reasoning and scientific discovery capabilities by developing theoretically complex and quantitatively challenging scientific equations in various fields of neuroscience, such as dynamical systems, nonlinear dynamical systems, dynamical systems on differential manifolds, neural field theory, nonlinear divergence theorems, nonlinear heat conduction equations and Laplace equations and their extensions on differential manifolds, nonlinear statistical analysis methods, deep learning, and other topics involving multiple fields. The current large language models may illustrate a certain degree of general intelligence, even if fundamentally it may be different from human intelligence. However, it does not mean the current LLMs can fully apply such ability in practical applications or that their reasoning potential can be fully tapped. It is essential to explore specific pathways and methods to cultivate their potential for scientific discovery. To accomplish this, we consider how to integrate them with common search engines (such as Google) capabilities and ChatGPT4o cross-modal abilities to better understand new disciplines and scientific discoveries. To this point, the major shortcoming of ChatGPTo1 is that it is not an end-to-end scientific discovery method and lacks the ability to achieve full automation. It also lacks methods for image analysis and full-scale data analysis, making it difficult to use simulation and data analysis to evaluate and test proposed new theories and methods.

## Introduction

The current developments in artificial intelligence (AI) increase the hope that AI can provide a powerful tool to facilitate scientific discovery and to generate and validate new ideas for scientific research autonomously (Si et al. 2024). Large Language Models (LLMs), such as GPT-4, have demonstrated remarkable capabilities in understanding and generating human-like text. Their potential extends beyond simple language tasks, offering transformative possibilities in scientific research of all fields. By leveraging vast amounts of data and advanced computational power, LLMs can assist researchers in generating novel ideas, automating routine tasks, and fostering interdisciplinary collaborations. These potentials are summarized as follows.

1. Efficient data mining and trend analysis. LLMs can efficiently extract key concepts, methodologies, and findings, as well as identify emerging trends. By drawing connections between disparate fields, LLMs can develop innovative transdisciplinary approaches by combining insights from various disciplines. They assist in organizing complex information into coherent structures (Lee et al. 2020; Wang et al. 2019).
2. Generating novel research ideas and hypotheses. Under appropriate prompts, the LLM can generate new concepts, identify new scientific research questions, and provide solutions to complex problems. They can generate plausible hypotheses based on existing data for researchers to conduct further empirical investigations. LLMs can predict potential outcomes of experiments or interventions, guiding experimental design (Surden 2019; Cohen 2015).
3. Automating scientific tasks. The LLM can automatically perform complex data analysis and interpretation, assisting in identifying optimal experimental conditions through predictive modeling (Jumper et al. 2021; Esteva et al. 2017).
4. Integration with domain-specific models. The LLMs help in developing domain-specific scientific models and extending the models across domains. By integrating LLMs with other AI methodologies (e.g., symbolic reasoning), we can generate more intelligent ideas and robust problem-solving based on mathematical and statistical models (Delépine et al. 2022).
5. On September 12, 2024, OpenAI released their new generative artificial intelligence system called ChatGPTo1. This new AI system, built upon “chain-of-thought” and reinforcement learning, has greatly enhanced logical reasoning abilities and can effectively solve various complex problems from elementary-level mathematical problems to modern scientific research issues in physics, chemistry, and biology. The new AI system can also combine reasoning with Internet search, significantly increasing ChatGPTo1’s knowledge and improving its performance on intellectual tasks.

The core of scientific research and invention is to discover previously unknown natural laws, propose new concepts, new hypotheses, theorems, new theories, data analysis methods, and new problem-solving approaches, as well as develop new products. We specifically tested ChatGPTo1’s existing reasoning and scientific discovery capabilities by studying complex and challenging scientific equations in the field of neuroscience, such as dynamical systems, nonlinear dynamical systems, dynamical systems on differential manifolds, neural field theory, nonlinear divergence theorems, nonlinear heat conduction equations and Laplace equations and their extensions on differential manifolds, nonlinear statistical analysis methods, deep learning, and other topics involving multiple fields. All previous large language models are pre-trained, meaning their logical reasoning and data analysis abilities are obtained through training on actual data. However, ChatGPTo1’s logical reasoning ability and capacity to discover new scientific ideas do not rely on data pre-training but are mainly acquired through chain-of-thought and reinforcement learning. While large language models based on pre-training may possess strong model intelligence, this intelligence is insufficient for discovering new scientific ideas and methods. On the contrary, ChatGPTo1 is more capable of developing novel ideas and discovering innovative methods. Therefore, as a reflection, models proposed in our series of articles on theoretical neuroscience research are new and difficult to find in previous literature. The large language models may have illustrated model performance that reflects a certain degree of “general intelligence”, even if fundamentally it may be different from human intelligence. However, it does not mean the current LLMs can fully apply such ability in practical applications or that their reasoning potential can be fully tapped. It is essential to explore specific pathways and methods to cultivate their potential for scientific discovery. To accomplish this, we consider how to integrate them with common search engines’ (such as Google’s) capabilities and ChatGPT4o’s cross-modal abilities to better understand new disciplines and scientific discoveries. To this point, the major shortcoming of ChatGPTo1 is that it is not an end-to-end scientific discovery method and lacks the ability to achieve full automation. It also lacks methods for image analysis and full-scale data analysis, making it difficult to use simulation and data analysis to evaluate and test proposed new theories and methods. Now we use several dynamic models of neural population dynamics as examples to illustrate the processes of using ChatGPTo1 for scientific discovery.

**Understanding Linear Dynamic Models of Neural Population Dynamics**

**Prompt: What is a linear dynamic model of neural population dynamics?**

**ChatGPTo1:**

**Understanding Linear Dynamic Models of Neural Population Dynamics**

## Introduction

Neural population dynamics studies how groups of neurons interact and evolve over time to produce complex brain functions such as perception, movement, and cognition. Instead of focusing on individual neurons, these models consider the average activity of large neural populations, making the analysis more tractable. Neuronal activity refers to the electrical and chemical signals that are generated and transmitted by neurons in the brain. It is often action potentials fired by a neuron or membrane potential of neurons (Figure 1). A **linear dynamic model** in this context refers to a mathematical framework where the change in neural activity is described using linear equations.

**Figure 1.**
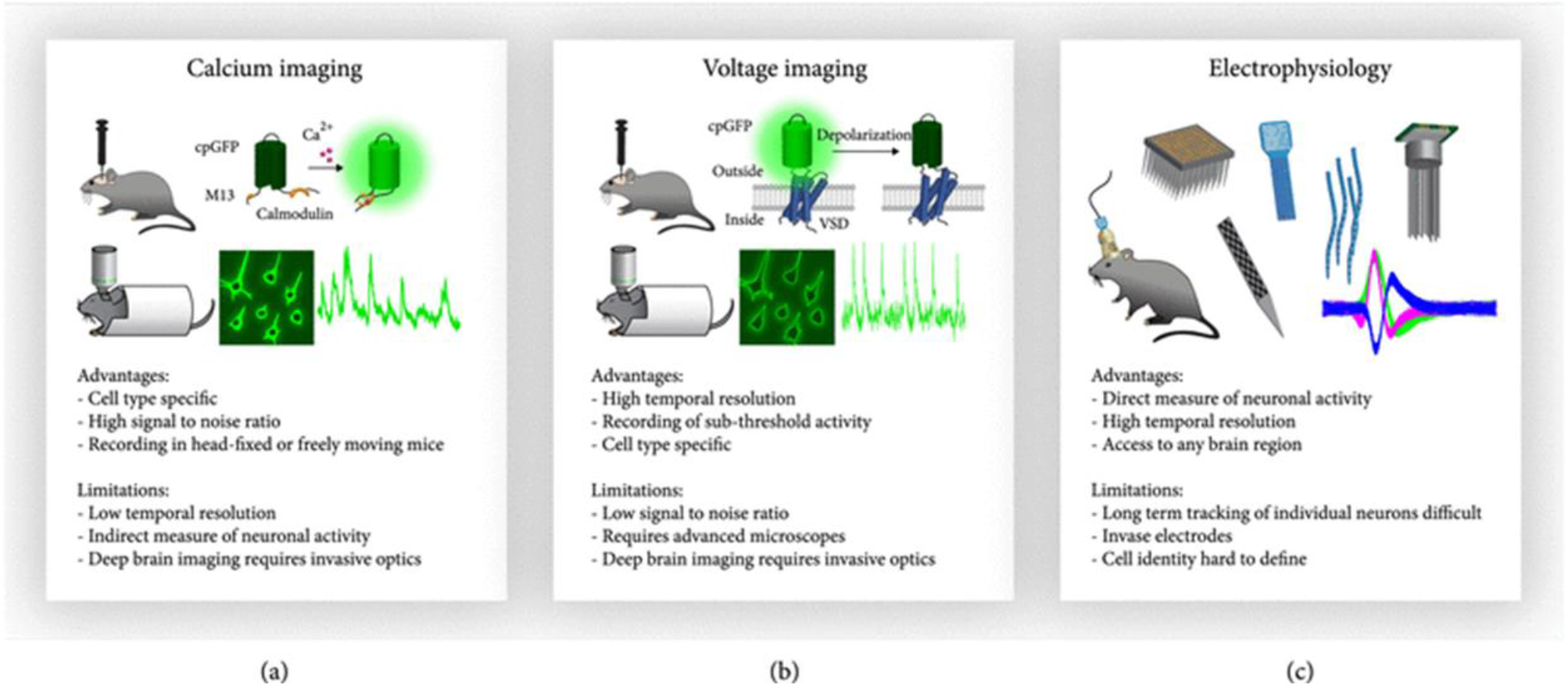
From Neurons to Cognition: Technologies for Precise Recording of Neural Activity Underlying Behavior.

### Linear Dynamic Models Explained

A linear dynamic model uses linear differential or difference equations to represent how neural populations change over time. The key feature of these models is that the rate of change of neural activity is a linear function of the current state and inputs. This means that the principles of superposition and homogeneity apply, simplifying analysis and solution.

### Mathematical Formulation

A general form of a linear dynamic model for neural population activity *u*(*x, t*) can be expressed as:

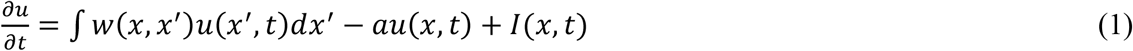

Where:

- *u*(*x, t*): Neural activity at position *x* and time *t*.
- *w*(*x, x*^′^): Linear connectivity kernel representing the influence of activity at *x*′ on *x*.
- *a*: Decay rate constant.
- *I*(*x, t*): External input to the neural population at position *x* and time *t*.

### Simplified Model Using Matrix Notation

In a discretized form for *N* neural populations, the model becomes:

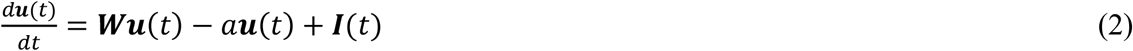

Where:

- ***u***(*t*): Vector of neural activities for each population at time *t*.
- ***W***: Connectivity matrix (linear operator).
- ***I***(*t*): Vector of external inputs.

**Prompt**

**In the linear dynamic model, please also present observation equation.**

**ChatGPTo1:**

**Observation Equation in Linear Dynamic Models of Neural Population Dynamics**

## Introduction

In linear dynamic models of neural population dynamics, we describe how neural activity evolves over time using linear differential or difference equations (the **state equations**). EEG, MEG and fMRI signals are important functional neuroimaging modalities for studying the temporal dynamics of neural activities and interactions with the accurate localization of neural activities. To connect these internal neural states to measurable data (such as EEG, MEG, fMRI signals, or behavioral outputs), we introduce the **observation equation**. The observation equation relates the hidden neural states to observable quantities, allowing us to interpret experimental data within the context of the model (Shenoy and Kao, 2021).

(Search Google for “Linear Dynamic Models of Neural Population Dynamics”)

### State Equation (System Model)

First, let’s recall the general form of the linear state equation for neural population dynamics:

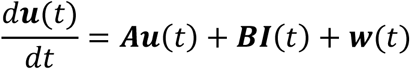

- ***u***(*t*): State vector representing neural activity of different populations at time *t*.
- ***A***: State transition matrix (connectivity matrix), defining interactions between neural populations.
- ***B***: Input matrix, mapping external inputs to the state space.
- ***I***(*t*): External input vector at time *t*.
- ***w***(*t*): Process noise vector, accounting for unmodeled dynamics or random fluctuations (typically assumed to be Gaussian with zero mean).

### Observation Equation (Measurement Model)

The **observation equation** links the internal neural states to the observable outputs:

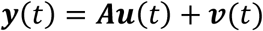

Where:

- ***y***(*t*): Observation vector at time *t*, containing measurable data.
- ***C***: Observation matrix, mapping the neural state space to the observation space.
- ***v***(*t*): Measurement noise vector, representing errors or uncertainties in the observations (typically Gaussian with zero mean).

### Explanation of Components

- **Observation Matrix *C*:** This matrix defines how each neural population contributes to the observed measurements. Each element *c*_*ij*_ represents the influence of the *j*-th neural population on the *i*-th observation channel.
- **Measurement Noise *v***(*t*): Accounts for inaccuracies in measurements due to sensor noise, environmental interference, or other factors not captured by the model.

### Role of the Observation Equation

- **Connecting Theory and Data:** It allows us to compare model predictions with actual experimental data, facilitating validation and refinement of the model.
- **State Estimation:** Essential for techniques like the Kalman filter, which estimate the hidden neural states ***u***(*t*) based on noisy observations ***y***(*t*).
- **Parameter Identification:** Helps in estimating model parameters (e.g., elements of ***A, B, C***) from observed data, improving the model’s accuracy.
- **Prediction and Control:** Enables forecasting future observations and designing interventions or control strategies.

### Example

#### Simple Neural Population Model

Consider a model with two neural populations:

1. **Excitatory population** (*u*_*E*_(*t*))
2. **Inhibitory population** (*u*_*I*_(*t*))

(Search google for “excitatory and inhibitory populations”)

Excitatory neurons are almost exclusively pyramidal cells, while inhibitory neurons form a diverse class with multiple subtypes. In many cases, inhibitory neurons exhibit greater firing rates than excitatory neurons, more attentional modulation, and a lesser degree of burstiness (Bittner et al. 2017)

#### State Equations

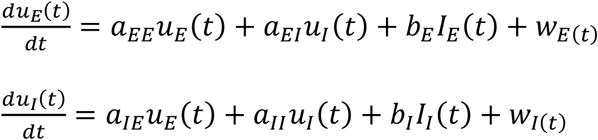

#### Observation Equation

Suppose we measure an electrical signal y(t)y(t)y(t) that is a linear combination of the activities of both populations:

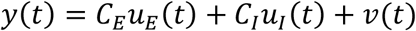

- *C*_*E*_, *C*_*I*_: Coefficients representing the contribution of excitatory and inhibitory populations to the observed signal.
- *v*(*t*):Measurement noise (e.g., sensor noise).

#### Matrix Formulation

- **State Vector:** 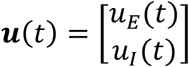
- **Observation Vector: *y***(*t*)=*y*(*t*)
- **Observation Matrix:** *C*= [*C*_*E*_ *C*_*I*_]

#### Interpretation in Neural Population Dynamics

- **EEG/MEG Measurements:** The observation equation models how electrical activity from neural populations contributes to scalp recordings.
- **fMRI Signals:** Relates neural activity to BOLD signals through hemodynamic response functions included in *C*.
- **Behavioral Outputs:** Connects neural states to observable behaviors or responses in tasks.

Assumed True Parameter Values:

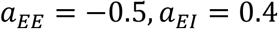

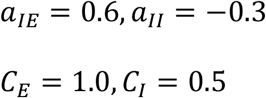

Initial Conditions:

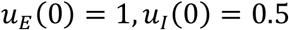

Time Discretization:

- Time step: Δ*t*= 0.1
- Time points: *t*= 0, 0.1, 0.2,…, 10 (Total of 101 time points)

Simulating the Electrical Signal as shown in Figure 4.

**Figure 2.**
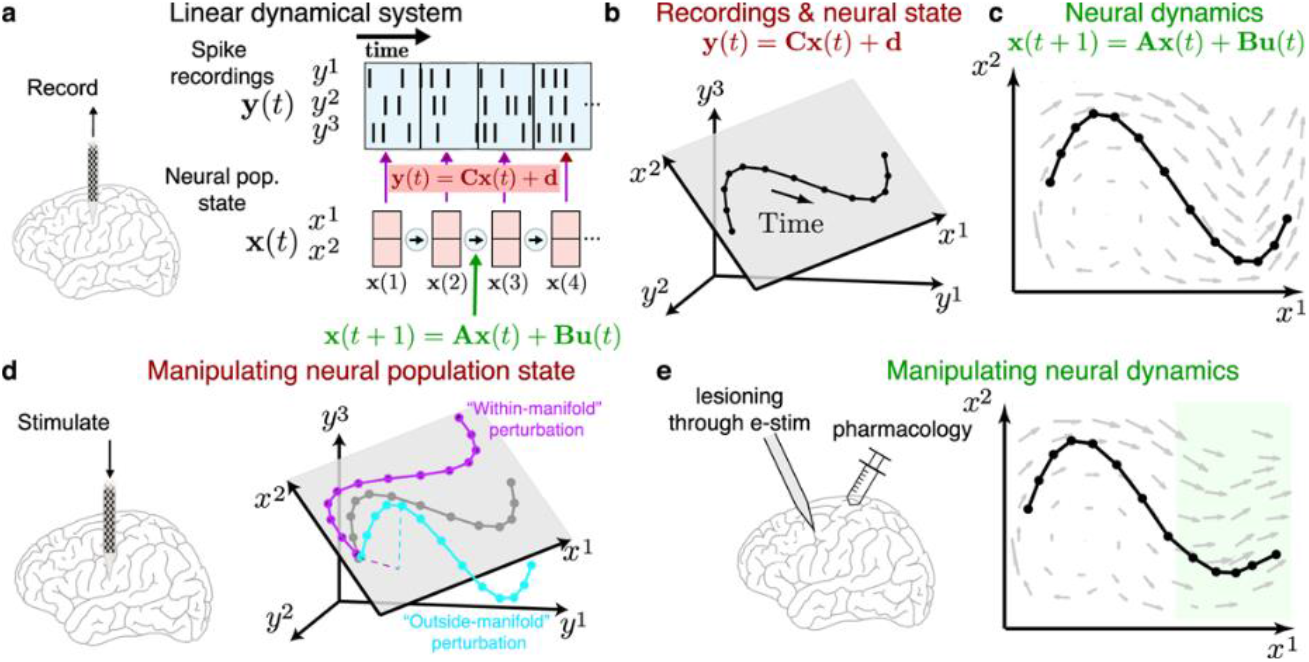
Overview of neural dynamics and manipulations (Shenoy and Kao, 2021)

**Figure 3.**
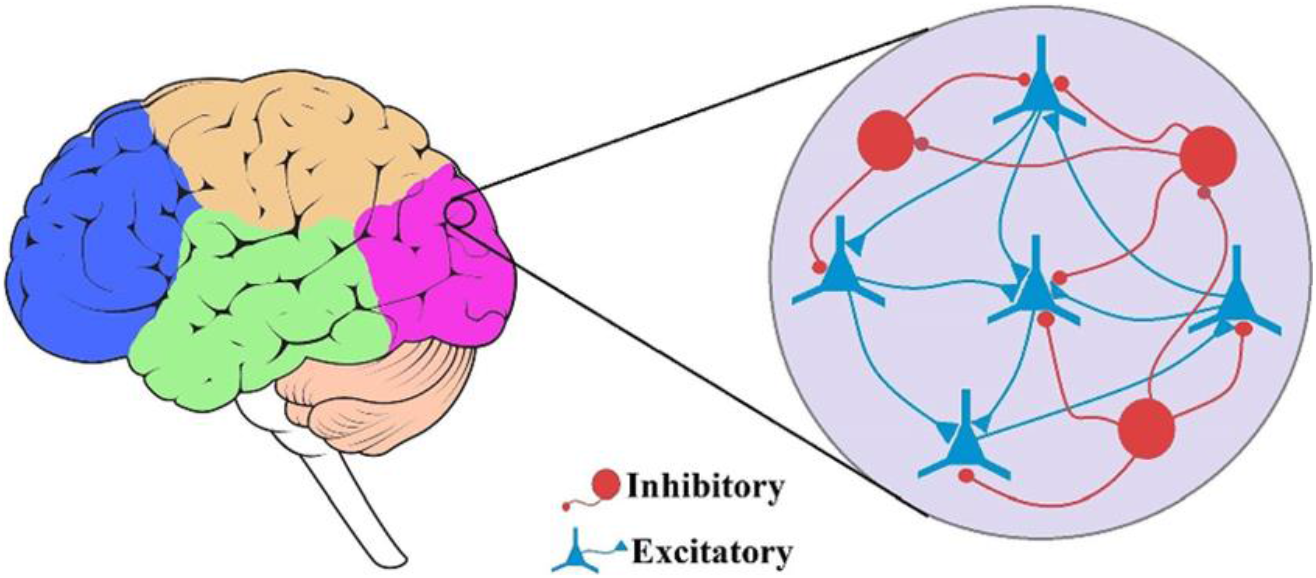
Connections of the excitatory (blue) and inhibitory (red) neurons in the cerebral cortex. (from online)

**Figure 4.**
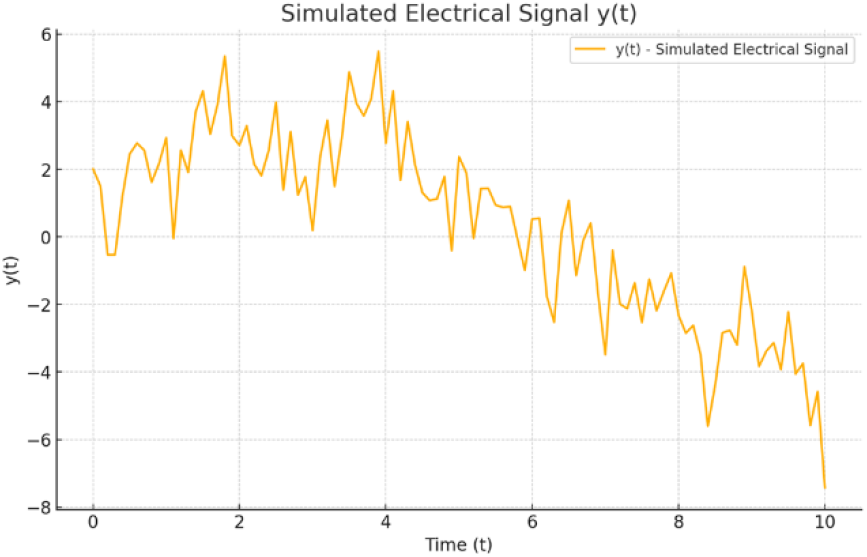
the plot of the simulated electrical signal *y*(*t*)

Here is the plot of the simulated electrical signal *y*(*t*) based on the linear dynamic model you described, with the specified parameter values and noise. The signal is generated over 101 time points in the interval [0,1]. Let me know if you’d like to =0.4 further analyze the data or explore another aspect of the mode.

**Multi-Area Neural Population Dynamics: Modeling Upstream to Downstream Brain Region Interactions**

**Prompt:**

**Please present and explain multi-area neural population dynamics model from upstream brain regions to downstream brain regions**

**ChatGPTo1**

**BEGIN**

Thought for a couple of seconds

Mapping neural dynamics

I’m thinking through how neural activity travels from upstream to downstream, detailing the process and its mathematical modeling.

**END**

## Introduction

The brain is a highly interconnected network composed of numerous regions that communicate to process information, generate behavior, and maintain homeostasis. Understanding how neural activity propagates across multiple brain areas is crucial for deciphering complex cognitive functions. Multi-area neural population dynamics models aim to capture these interactions by representing the collective behavior of neuronal populations in different brain regions and modeling the pathways through which they influence each other.

### (1). Overview of Neural Population Dynamics

**Neural Population Dynamics** refers to the study of how groups of neurons behave collectively over time. Instead of focusing on individual neurons, these models consider the average activity of large ensembles, which simplifies analysis and highlights emergent properties not evident at the single-neuron level.

### (2). Multi-Area Models: Connecting Brain Regions

**Multi-area models** extend single-area neural population models by incorporating multiple interconnected brain regions. Each area is represented by its own population dynamics, and the connections between areas are modeled to capture the flow of neural activity from upstream to downstream regions.

**Key Components:**

1. **Neural Populations:** Each brain area is represented by excitatory and inhibitory populations.
2. **Local Dynamics:** Within each area, neurons interact through local connectivity.
3. **Inter-Area Connectivity:** Connections between different areas represent the pathways through which activity propagates.

### (3). Mathematical Formulation

#### (3.1) State Variables

Let:

- *u*_*i*_(*t*): Neural activity vector of area *i* at time *t*, comprising excitatory (*u*_*E,i*_(*t*)) and inhibitory (*u*_*I,i*_(*t*)) populations.

#### (3.2) State Equations

The dynamics of each area *i* are governed by differential equations that include:

- **Intrinsic Dynamics:** Local interactions within area *i*
- **Input from Other Areas:** Influences from upstream areas *j*.

##### General Form

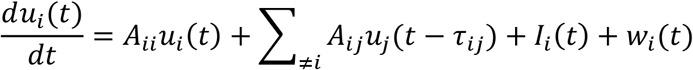

where

- *A*_*ii*_: Local connectivity matrix of area *i*.
- *A*_*ij*_: Connectivity matrix from area *j* to area *i*.
- *τ*_*ij*_: Time delay in transmission from area *j* to area *i*.
- *I*_*i*_(*t*): External input to area *i*.
- *w*_*i*_(*t*): Process noise.

#### (3.3) Inter-Area Connectivity

- **Feedforward Connections:** From upstream to downstream areas (e.g., *A*_*ij*_where *j*is upstream of *i*).
- **Feedback Connections:** From downstream back to upstream areas (e.g., *A*_*ji*_).

**Example:**

For two areas, upstream (area 1) and downstream (area 2):

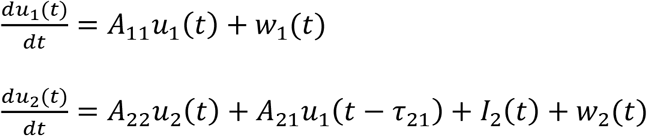

- **Upstream Area (1):** Affects downstream area through *A*_21_*u*_1_(*t − τ*_21_).
- **Downstream Area (2):** Receives input from area 1.

### (4). Detailed Explanation

#### (4.1) Local Dynamics within Each Area

Each area has its own excitatory and inhibitory populations interacting locally.

##### State Equations for Area iii

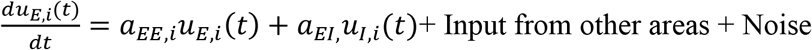

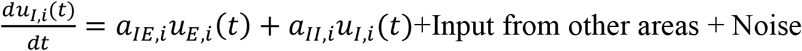

- *a*_*EE,i*_, *a*_*EI,i*_, *a*_*IE,i*_*a*_*II,i*_Local connectivity parameters within area *i*.

#### (4.2) Inputs from Other Areas

- **Feedforward Input:** Activity from upstream areas influences the excitatory and inhibitory populations of the downstream area.
- **Inclusion of Time Delays** *τ*_*ij*_: Accounts for the finite speed of sigjnal propagation along axonal pathways.

##### Input Term from Area *j* to Area *i*

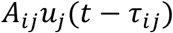

***A***_***i****j*_: Connectivity matrix specifying how populations in area Type equation here. affect those in area *i*.

#### (4.3) Example: Three-Area Model

Consider three areas:

1. **Area V1 (Primary Visual Cortex):** Upstream area processing basic visual features.
2. **Area V2:** Intermediate processing area
3. **Area V4:** Downstream area involved in higher-level visual processing.

##### State Equations

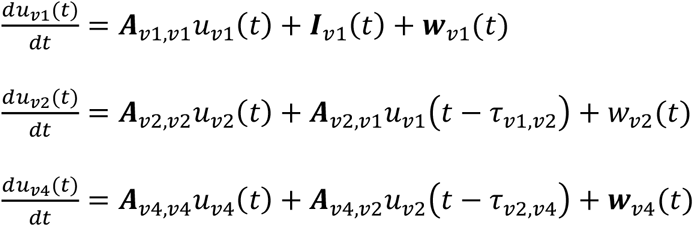

### (5). Observation Equations

To connect the model to measurable data (e.g., EEG, fMRI), observation equations are used:

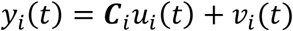

- *y*_*i*_(*t*): Observed signal from area iii.
- ***C***_***i***_: Observation matrix mapping neural activity to measured signals.
- *v*_*i*_(*t*): Measurement noise.

### (6). Analysis and Simulation

#### (6.1) Numerical Simulation

- **Initial Conditions:** Set initial neural activities for each area.
- **Time Integration:** Use numerical methods (e.g., Euler, Runge-Kutta) to simulate the model over time.
- **Incorporate Noise:** Add process and measurement noise to reflect biological variability.

#### (6.2) Parameter Estimation

- **Data Fitting:** Adjust model parameters to fit experimental data.
- **Techniques Used:**
  ∘ Least squares optimization.
  ∘ Maximum likelihood estimation.
  ∘ Bayesian inference.

#### (6.3) Stability and Dynamics

- **Eigenvalue Analysis:** Study the stability of each area by analyzing the eigenvalues of the local connectivity matrices.
- **Propagation of Activity:** Examine how perturbations in upstream areas affect downstream regions.
- **Oscillations and Rhythms:** Investigate how inter-area interactions contribute to neural oscillations observed in brain signals.

### (7). Biological Interpretations

#### (7.1) Functional Connectivity

- **Directional Influence:** The model captures the directionality of neural interactions from upstream to downstream regions.
- **Information Processing:** Demonstrates how sensory inputs are transformed and integrated across brain areas.

#### (7.2) Pathological Conditions

- **Disrupted Connectivity:** Alterations in inter-area connections can model pathological conditions (e.g., schizophrenia, epilepsy).
- **Intervention Effects:** Simulate the impact of interventions (e.g., pharmacological agents, brain stimulation) on neural dynamics.

### (8.) Example Application

#### (8.1) Visual Processing Pathway

- **Objective:** Model how visual information flows from the retina through the visual cortex.
- **Areas Involved:** Retina → LGN (Lateral Geniculate Nucleus) → V1 → Higher Visual Areas.
- **Model Setup:**
  ∘ **Retinal Input** *I*_*retina*_(*t*): External stimulus.
  ∘ **Connectivity Matrices:** Define how each area influences the next.
  ∘ **Time Delays:** Account for synaptic and transmission delays.

#### (8.2) Simulation Results

- **Stimulus Response:** Observe how a visual stimulus leads to activity changes across areas.
- **Latency Analysis:** Measure the time it takes for activity to propagate downstream.
- **Amplitude Modulation:** Examine how the strength of upstream activity affects downstream responses.

### (9.) Extensions of the Model (9.1) Nonlinear Dynamics

- Incorporate nonlinear functions (e.g., sigmoid activation) to model neuronal firing rates more accurately.

#### (9.2) Plasticity

- Include mechanisms for synaptic plasticity to model learning and adaptation.

#### (9.3) Network Topology

- Expand the model to include more areas and complex connectivity patterns (e.g., recurrent loops, parallel pathways).

### (10.) Summary

**Multi-area neural population dynamics models** provide a powerful framework for understanding how neural activity propagates from upstream to downstream brain regions. By representing each area with its own population dynamics and modeling the connections between areas, these models capture the complex interactions underlying brain function.

**Key Takeaways:**

- **Integration of Multiple Scales:** Bridges the gap between local neural circuits and large-scale brain networks.
- **Versatility:** Applicable to various sensory modalities and cognitive processes.
- **Research Utility:** Aids in interpreting experimental data, testing hypotheses about brain function, and developing treatments for neurological disorders.

**Incorporating Synaptic Plasticity into Multi-Area Neural Population Dynamics Models**

**Prompt**

**Please incorporate mechanisms for synaptic plasticity to model learning and adaptation into the above models**.

**ChatGPTo1**

**Thought for 6 seconds**

## Introduction

In the previous discussion, we explored multi-area neural population dynamics models that describe how neural activity propagates from upstream to downstream brain regions. These models capture the interactions between excitatory and inhibitory populations within each area and the connections between different areas. However, to model learning and adaptation, it’s essential to incorporate mechanisms of **synaptic plasticity**—the ability of synapses to strengthen or weaken over time in response to activity levels.

(Search Google for “synaptic plasticity to model learning and adaptation”)

(Neural plasticity refers to the possibility of altering the strength of connections within the nervous system through experience or injury. Plasticity can act to reorganize either the structure or the function of neurons and is necessary not only for neural networks to acquire new capabilities, but also for them to remain robust and stable over time (Appelbaum et al. 2023). Synaptic plasticity refers to the adaptive changes that occur at the synapse, resulting in the strengthening or weakening of synaptic connections. It is a fundamental mechanism involved in learning and memory, and it is influenced by factors such as neurotransmitter release and activation of neighboring structures (Weishaupt 2017).)

### (1.) Synaptic Plasticity: Mechanisms of Learning and Adaptation

**Synaptic Plasticity** is the process by which the efficacy of synaptic transmission is modified, enabling the brain to adapt to new information, store memories, and refine neural circuitry. Key mechanisms include:

- **Hebbian Plasticity**: Synapses are strengthened when the presynaptic and postsynaptic neurons have correlated activity (“cells that fire together wire together”). ((Google search for “Hebbian plasticity”)
- **Spike-Timing-Dependent Plasticity (STDP)**: Synaptic changes depend on the precise timing of spikes between presynaptic and postsynaptic neurons. ((Search google for “Spike-Timing-Dependent Plasticity (STDP)”) The relative magnitude (y-axis, in %) and spike-time intervals (temporal windows, x-axis, in ms) are simplified and presented schematically and arbitrary in A–C. A In the classical concept, STDP is equally balanced for t-LTD and t-LTP: postsynaptic spiking occurring before presynaptic activity drives t-LTD (in red, pre-post timing interval: Δt < 0) whereas presynaptic spiking occurring before postsynaptic spike leads to t-LTP (in green, pre-post timing interval: Δt > 0). B Schematic showing that the sign of STDP is under the control of external input at CA3-CA1 synapses from 13–18 postnatal days (P13–P18) mice. C Dopamine (DA) influences the sign and the time window for STDP in CA3-CA1 synapses at P13–P18. D STDP time window in CA3-CA1 synapses changes during development. A post-pre protocol (Δt from *−*35 to *−*18 ms) induces t-LTD at P13– P21 (in red). magnitudes that may follow this behavior.
- **Homeostatic Plasticity**: Mechanisms that maintain overall neural activity within functional ranges.
  - **((Search Google for “Homeostatic Plasticity”)**

Incorporating these mechanisms into neural population models allows us to simulate how neural circuits learn and adapt over time.

### (2.) Modifying the Multi-Area Model to Include Synaptic Plasticity

#### (2.1) State Variables and Synaptic Weights

We introduce time-dependent synaptic weights to represent the strength of connections both within and between areas:

- **Within-Area Synaptic Weights**: ***W***_*ii*_(*t*)
- **Between-Area Synaptic Weights**: ***W***_*ij*_(*t*)

These weights evolve over time according to plasticity rules.

#### (2.2) Updated State Equations

The dynamics of neural activity *u*_*i*_(*t*) in area *i* are now influenced by the time-varying synaptic weights:

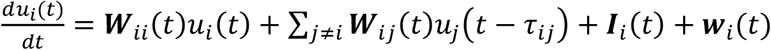

#### 2.3 Synaptic Plasticity Equations

The synaptic weights evolve according to plasticity rules, which are functions of the pre- and postsynaptic neural activities:

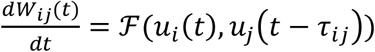

- ℱ: Function defining the plasticity rule.

### (3.) Hebbian Plasticity in Neural Population Models

#### 3.1 Hebbian Learning Rule

A simple Hebbian learning rule can be formulated as:

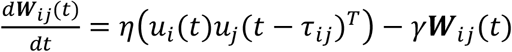

Where:

- *η*: Learning rate constant.
- *γ*: Weight decay constant (prevents unbounded growth of weights).
- *u*_*i*_(*t*) and*u*_*j*_(*t − τ*_*ij*_) : Pre- and postsynaptic activities.
- *T*: Transpose operation (if *u*_*i*_ and *u*_*j*_ are vectors).

#### (3.2) Incorporating Hebbian Plasticity into the Model

##### State Equations with Plasticity

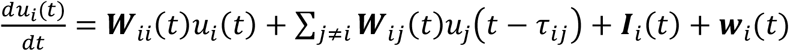

##### Synaptic Weight Equations

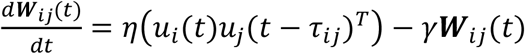

### (4.) Spike-Timing-Dependent Plasticity (STDP)

While STDP is typically formulated at the level of spiking neurons, we can approximate its effects in population models.

#### 4.1 STDP-Based Plasticity Rule

The change in synaptic weight depends on the timing difference between pre- and postsynaptic activities:

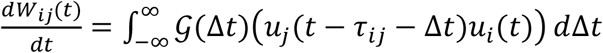

- 𝒢(Δ*t*)STDP learning window function, typically with exponential decay for positive and negative Δ*t*.

#### (4.2) Simplified STDP in Population Models

Assuming a continuous-time approximation:

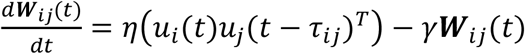

This resembles the Hebbian rule but can be adjusted to reflect the asymmetric learning window of STDP.

### (5.) Homeostatic Plasticity

To prevent runaway excitation or suppression, homeostatic mechanisms adjust synaptic strengths to maintain stable activity levels.

#### (5.1) Homeostatic Plasticity Rule

Synaptic weights are adjusted based on the deviation of neural activity from a target level *u*_*taregt*_:

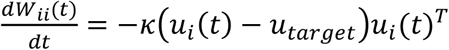

- *κ*: Homeostatic adjustment rate.

### (6.) Complete Model with Synaptic Plasticity

Combining the above mechanisms, the multi-area model becomes:

#### (6.1) State Equations

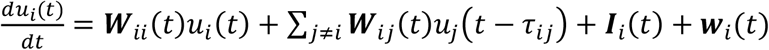

#### (6.2) Synaptic Weight Equations

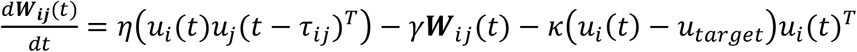

- **Within-Area Weights** (***W***_*ii*_(*t*)): Include both Hebbian and homeostatic plasticity.
- **Between-Area Weights** (***W***_*ij*_(*t*)): Primarily governed by Hebbian or STDP rules.

### (7.) Biological Interpretation

#### (7.1) Learning and Memory Formation

- **Strengthening of Synapses:** Repeated co-activation of neurons in different areas leads to strengthening of synaptic connections, encoding new information.
- **Propagation of Learning:** Changes in upstream areas affect downstream areas through modified synaptic weights, allowing for the transfer and consolidation of learning.

#### (7.2) Adaptation to Inputs

- **Experience-Dependent Plasticity:** The model adapts to external inputs *I*_*i*_(*t*), modifying synaptic strengths to optimize responses.
- **Critical Periods:** Time windows where plasticity is heightened, enabling rapid learning (can be modeled by adjusting *η*).

### (8.) Example Simulation

#### (8.1) Setup

- **Two Areas:** Area 1 (upstream) and Area 2 (downstream).
- **Initial Synaptic Weights:** Random small values.
- **External Input:** Repeated stimulus to Area 1.

#### (8.2) Simulation Steps

1. **Initialize Variables:**
  ∘ Set initial *u*_*i*_(0), ***W***_*ij*_(0), and parameters *η, γ, κ*
2. **Time Integration:**
  ∘ For each time step *t*:
    ▪ Update neural activities *u*_*i*_(*t*) using the state equations.
    ▪ Update synaptic weights ***W***_*ij*_(*t*) using the plasticity equations.
3. **Observation:**
  ∘ Monitor changes in synaptic weights and neural activities over time.

#### (8.3) Expected Outcomes

- **Synaptic Strengthening:** Synapses between co-active neurons in different areas become stronger.
- **Activity Patterns:** Neural activities in downstream areas become more responsive to inputs due to strengthened connections.
- **Stabilization:** Homeostatic plasticity prevents excessive excitation or inhibition. **((Search Google for “Homeostatic plasticity prevents excessive excitation or inhibition”)** The National Institute of Mental Health estimates 21.4% of U.S. adults experience a mood disorder, these include Major Depressive Disorder or Bipolar Disorder, at some point in their lives. Identification of rapid acting treatments with predictable outcomes for these disorders is an urgent societal need. Since the middle of the 20th century, neuropharmacological treatments primarily targeting monoaminergic neurotransmission have reached the clinic with varying degrees of efficacy for the treatment of certain mood disorders. The unexpected discovery that ketamine exerts rapid antidepressant action provides an opportunity to identify specific synaptic substrates that are key to triggering antidepressant effects, which may ultimately serve as new targets for drug development. Some forms of plasticity act in an adaptive fashion where changes in neuronal excitability or synaptic strength follow the direction of the manipulation and thus act to reinforce the demand imposed on the system via positive feedback regulation of neuronal or synaptic biophysical parameters (Figure 8).

**Figure 5.**
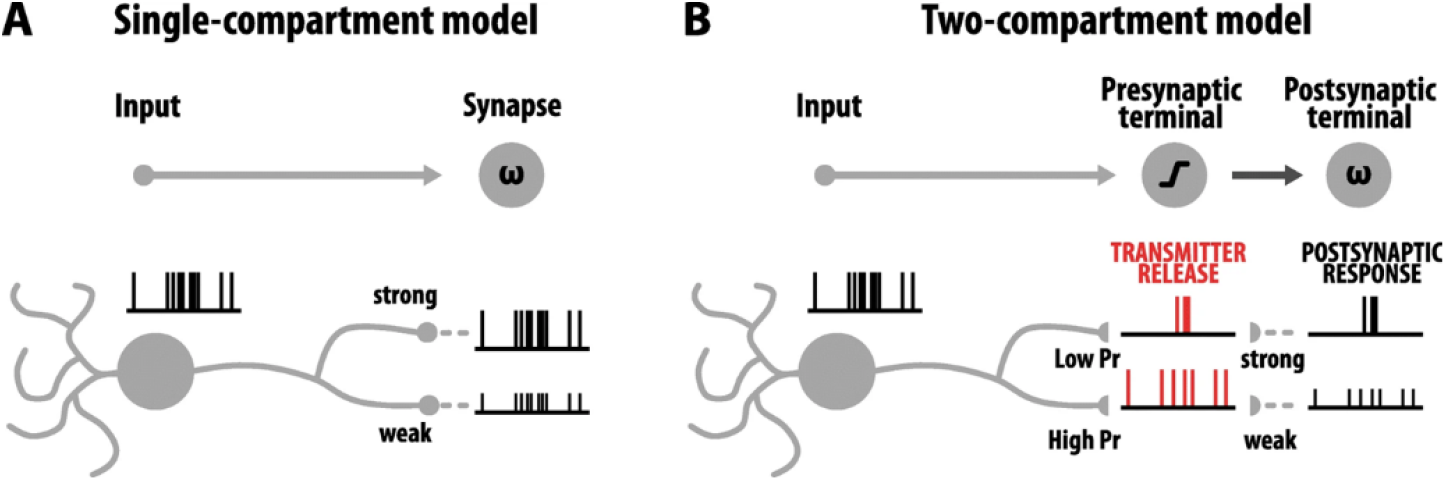
The two-compartment model of the synapse. a In the conventional single-compartment model, inputs are scaled linearly by the synapse, often characterised by a weighting factor w. b The addition of the presynaptic terminal results in a dynamic temporal filter prior to the postsynaptic gain control. This leads to a temporal decomposition of the presynaptic input spike train across presynaptic terminals along the axon. Importantly, this filter is dynamically regulated by ongoing neural activity to adjust to the statistics of pre- and postsynaptic activity. (Tong et al. 2020))

**Figure 6.**
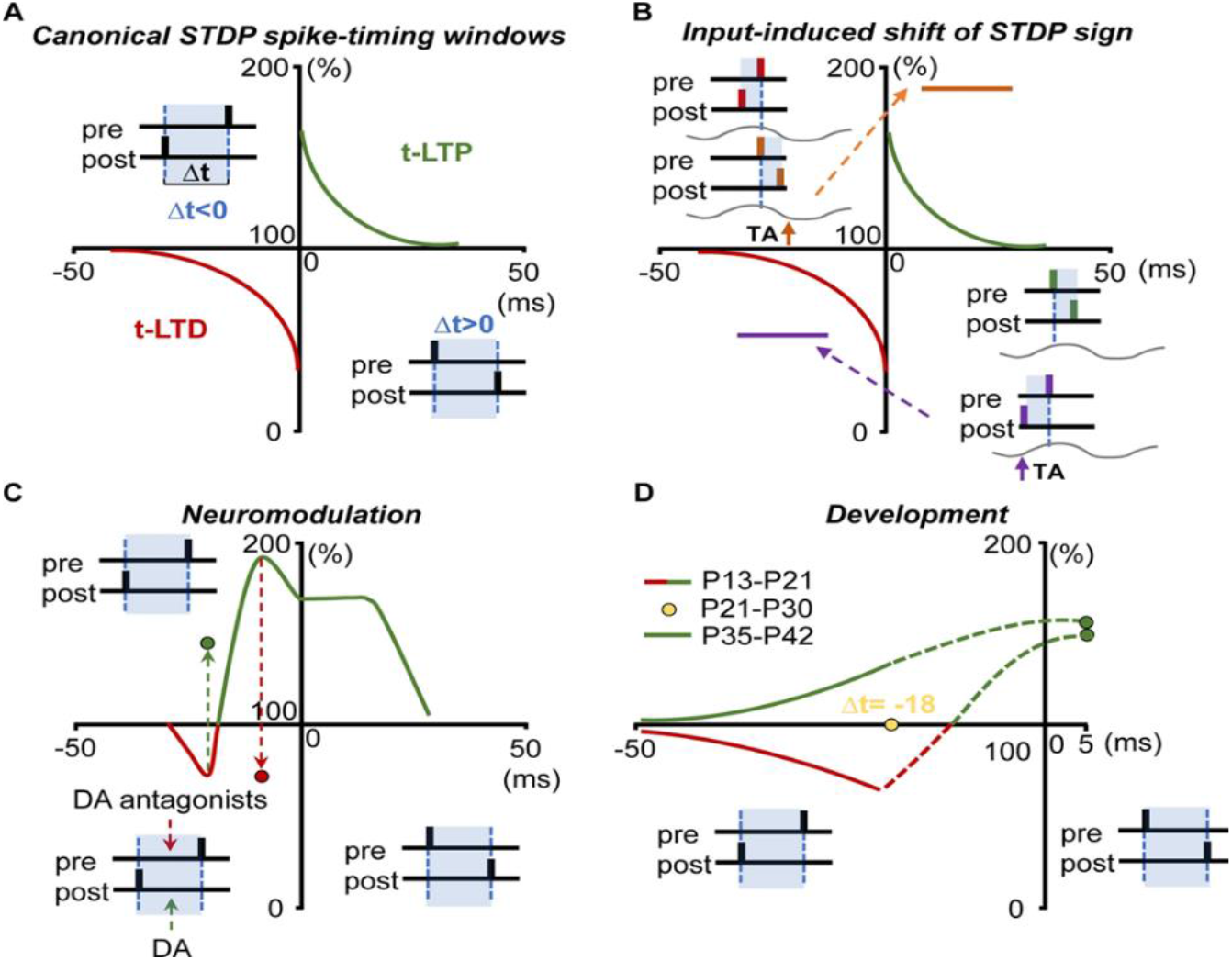
Schematic diagrams showing the diversity of STDP forms and some of the factors that might influence its expression, sign, and magnitude (Andrade-Talavera et al. 2023).)

**Figure 7.**
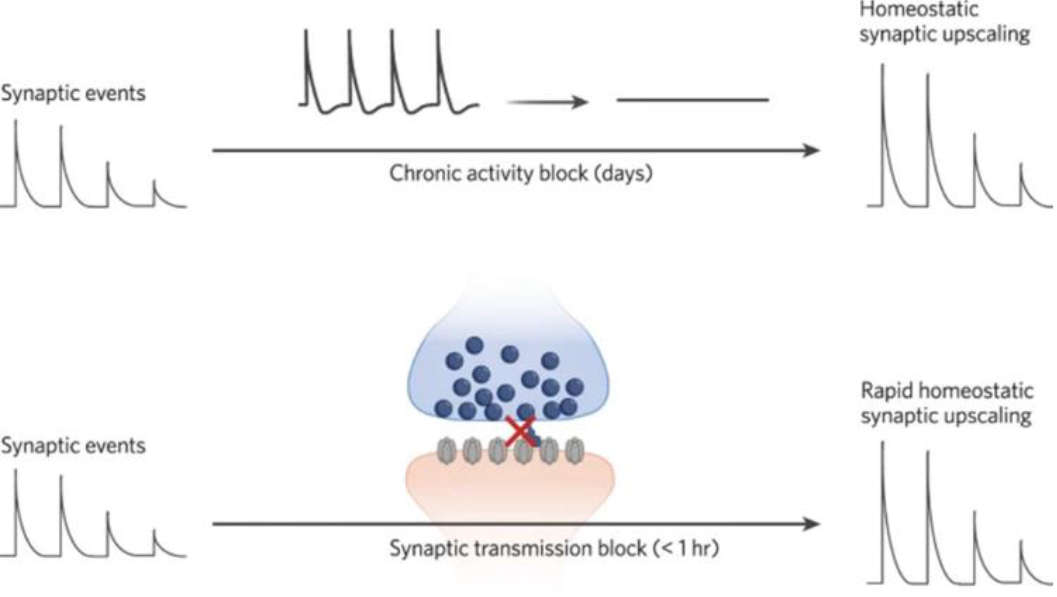
Rapid versus slow homeostatic plasticity. It depicts the impact of synaptic transmission blockade on the rapidity of homeostatic plasticity. While classical experiments relying on activity blockade (via blockade of voltage-gated Na+ channels or postsynaptic AMPA receptors) demonstrated a slow process that occurs over days, block of synaptic NMDA receptor facilitates plasticity down to hours or less (Kavalali and Monteggia, 2023).

**Figure 8.**
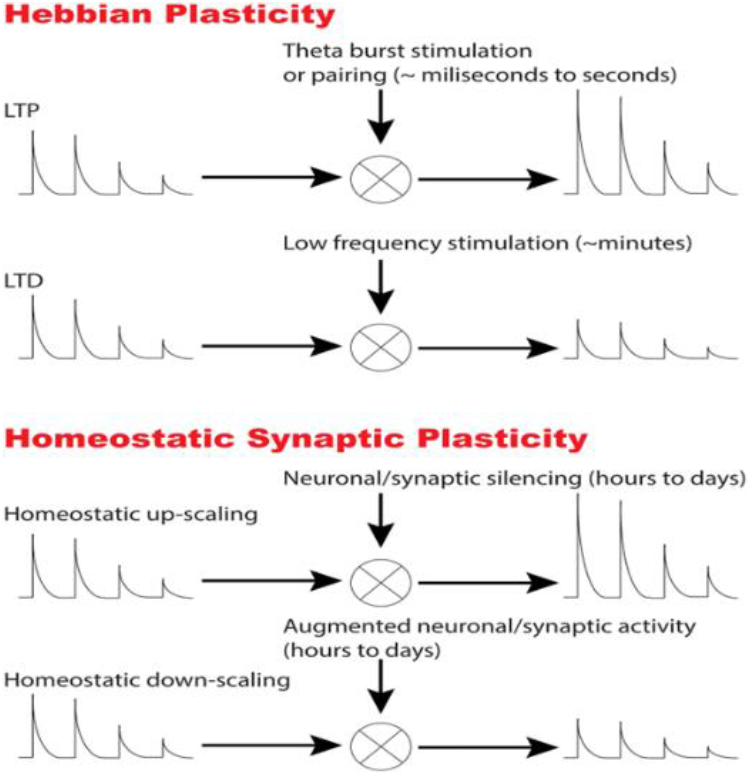
Hebbian versus Homeostatic Synaptic Plasticity This figure depicts key distinguishing properties of Hebbian versus Homeostatic synaptic plasticity. During Hebbian plasticity, synaptic strength is modified in the same direction of the applied stimuli. In contrast, homeostatic plasticity changes in neuronal network activity or alterations in synaptic inputs lead to an adjustment of synaptic strengths in the opposite direction (Kavalali et al. 2020).

### (9.) Analysis and Implications

#### (9.1) Network Connectivity Evolution

- **Structural Changes:** Synaptic plasticity leads to changes in the effective connectivity between brain regions.
- **Functional Connectivity:** Altered synaptic strengths reflect changes in functional connectivity observed in neuroimaging studies.

#### (9.2) Learning Dynamics

- **Skill Acquisition:** The model can simulate how repeated practice strengthens relevant neural pathways.
- **Memory Trace Formation:** Persistent changes in synaptic weights represent the formation of memory traces.

### (10.) Extensions and Considerations

#### (10.1) Nonlinear Neural Dynamics

- Incorporate nonlinear activation functions (e.g., sigmoid) to model saturation effects in neuronal firing rates.

#### (10.2) Spike-Based Models

- Transition to spiking neuron models for finer temporal resolution and more accurate STDP implementation.

#### (10.3) Neuromodulation Effects

- Include the influence of neuromodulators (e.g., dopamine) on plasticity rules to simulate reward-based learning.

#### (11.) Summary

Incorporating synaptic plasticity into multi-area neural population dynamics models enriches their ability to simulate learning and adaptation processes in the brain. By allowing synaptic weights to evolve based on neural activity, these models can capture the dynamic nature of neural circuits as they respond to experiences and stimuli.

**Incorporating Synaptic Plasticity Modeled by Kolmogorov-Arnold Networks into Multi-Area Neural Population Dynamics Models**

**Prompt**

**Please Incorporate Synaptic Plasticity that is modeled by Kolmogorov-Arnold Networks into Multi-Area Neural Population Dynamics Models**.

**ChatGPTo1**

**Thought for 11 seconds BEGIN**

## Understanding Kolmogorov-Arnold Networks

To address the user’s request, I’m tasked with defining Kolmogorov-Arnold networks, explaining their role in synaptic plasticity, and integrating this into multi-area neural population dynamics models.

## Understanding synaptic plasticity

I’m digging into how synaptic plasticity is modeled, emphasizing accurate factual claims to avoid errors.

## Understanding Kolmogorov-Arnold representation

I’m thinking through how the Kolmogorov-Arnold representation theorem applies to neural networks, suggesting they can approximate any continuous function by combining continuous functions of one variable.

## Integrating novel elements

I’m mapping out how synaptic plasticity could be modeled using Kolmogorov-Arnold networks and how to incorporate this into multi-area neural population dynamics models.

## Mapping out the model

I’m focusing on how Kolmogorov-Arnold networks can approximate functions and model synaptic plasticity, integrating them into neural population dynamics models.

**END**

## Introduction

Synaptic plasticity is a fundamental mechanism underlying learning and memory in the brain. Modeling synaptic plasticity within neural population dynamics provides insights into how neural circuits adapt and reorganize in response to stimuli. The **Kolmogorov-Arnold representation theorem** offers a mathematical framework for approximating multivariate functions using superpositions of univariate functions. **Kolmogorov-Arnold Networks (KANs)** leverage this theorem to approximate complex functions and can be applied to model synaptic plasticity in neural networks.

In this explanation, we will:

1. *Explain the Kolmogorov-Arnold representation theorem and Kolmogorov-Arnold Networks*.
2. *Describe how KANs can model synaptic plasticity*.
3. *Incorporate synaptic plasticity modeled by KANs into multi-area neural population dynamics models*.
4. *Discuss the implications and potential applications of this approach*.

### (1.) The Kolmogorov-Arnold Representation Theorem and Kolmogorov-Arnold Networks

#### (1.1) Kolmogorov-Arnold Representation Theorem

The **Kolmogorov-Arnold representation theorem** is a result in mathematical analysis concerning the representation of multivariate continuous functions. It states that any continuous function of several variables can be represented as a finite sum of continuous functions of one variable and addition.

##### Kolmogorov’s Theorem (1957)

For any continuous function *f*: *R*^*n*^ → *R*, there exist continuous functions ∅_*i*_ and *ψ* such that:

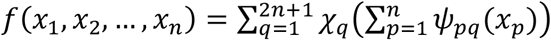

- *χ*_*q*_ and *ψ*_*pq*_are continuous univariate functions.

##### Implications

- **Function Approximation:** Any multivariate function can be decomposed into superpositions of univariate functions.
- **Dimensionality Reduction:** Simplifies the complexity of high-dimensional functions.

#### (1.2) Kolmogorov-Arnold Networks (KANs)

**Kolmogorov-Arnold Networks** are neural network architectures inspired by the representation theorem. They are designed to approximate multivariate functions using layers of neurons that implement univariate functions and summations.

##### Structure of KANs

1. **First Layer (Univariate Functions):** Applies continuous univariate functions *ψ*_*pq*_ to each input variable *χ*_*p*_.
2. **Summation Layer:** Sums the outputs of the first layer.
3. **Output Layer (Univariate Functions):** Applies continuous univariate functions *χ*_*q*_ to the summed inputs and sums the results to produce the output.

##### Properties

- **Universal Approximation:** KANs can approximate any continuous function on compact subsets of *R*^*n*^.
- **Layered Architecture:** Resembles feedforward neural networks with specific activation functions and connectivity.

### (2.) Modeling Synaptic Plasticity with Kolmogorov-Arnold Networks

#### (2.1) Synaptic Plasticity in Neural Networks

**Synaptic Plasticity:** The ability of synapses to strengthen or weaken over time in response to increases or decreases in their activity.

- **Hebbian Plasticity:** Synapses strengthen when pre- and postsynaptic neurons are co-active.
- **Spike-Timing-Dependent Plasticity (STDP):** Synaptic changes depend on the timing between pre- and postsynaptic spikes.

#### (2.2) Representing Synaptic Plasticity with KANs

##### Motivation

- **Complex Dependencies:** Synaptic changes may depend on multiple variables (e.g., pre- and postsynaptic activity, neuromodulators).
- **Function Approximation:** KANs can approximate the synaptic update rules as functions of relevant variables.

##### Modeling Approach

- **Synaptic Update Function:** Define the change in synaptic weight Δ*w*_*ij*_ as a function *f*(*u*_*i*_, *u*_*j*_, *θ*), where:
  ∘ *u*_*i*_: Activity of presynaptic neuron i.
  ∘ *u*_*j*_: Activity of postsynaptic neuron *j*.
  ∘ *θ*: Other variables (e.g., time, neuromodulators).
- **Approximation with KANs:** Use a Kolmogorov-Arnold Network to approximate *f*(*u*_*i*_, *u*_*j*_, *θ*).

##### Advantages

- **Flexibility:** Can model complex, nonlinear synaptic plasticity rules.
- **Dimensionality Reduction:** Simplifies high-dimensional dependencies into univariate functions.

### (3.) Incorporating KAN-Modeled Synaptic Plasticity into Multi-Area Neural Population Dynamics Models

#### (3.1) Multi-Area Neural Population Dynamics Models

**Overview**

- **Areas:** Multiple brain regions (areas) each containing neural populations.
- **State Variables:** *u*_*i*_(*t*) represents the neural activity in area *i* at time *t*.
- **Connectivity:** Synaptic weights ***W***_*ij*_(*t*) represent connections from area *j* to area *i*.

##### State Equations

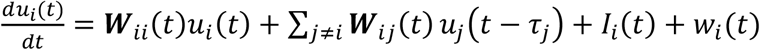

#### (3.2) Synaptic Plasticity Modeled by KANs

##### Synaptic Weight Dynamics

- **Synaptic Change:** Δ*W*_*ij*_(*t*)= *f*_*ij*_(*u*_*i*_(*t*), *u*_*j*_(*t − τ*_*j*_), *θ*_*ij*_)
- **Function Approximation:** Use KANs to approximate *f*_*ij*_.

##### Implementing KANs in Synaptic Updates

1. **Define Univariate Functions:**
  ∘ 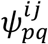 and 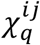 specific to synapses from area *j* to area *i*.
2. **Synaptic Update Rule** (Figure 9):

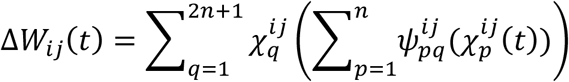

**Figure 9.**
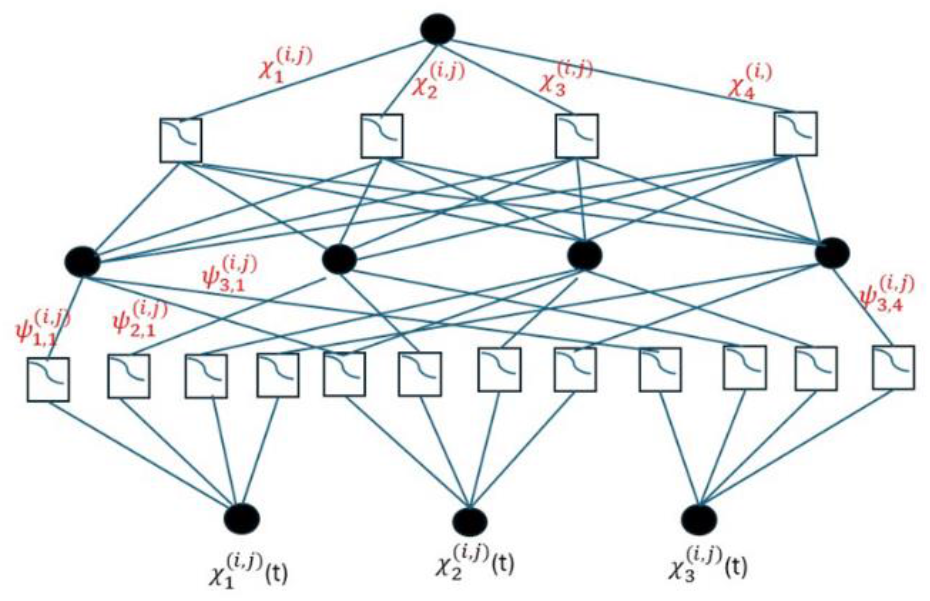
Synaptic Plasticity Modeled by KANs.

**Figure 10.**
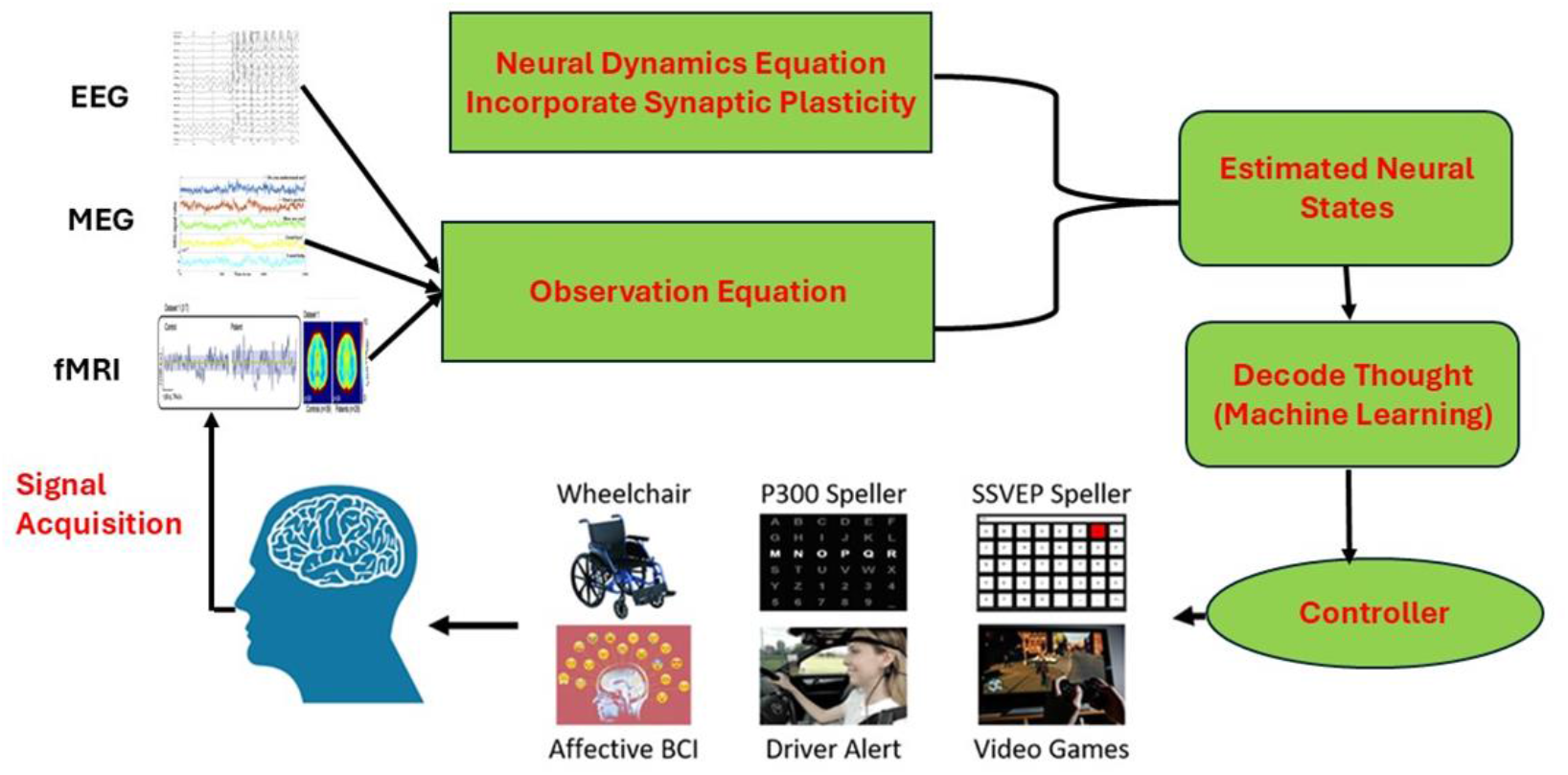
A diagram for describing dynamic population models acting on Brain-Computer Interface

where:

- 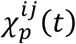 are variables influencing synaptic plasticity (e.g., *u*_*i*_(*t*), *u*_*j*_(*t − τ*_*ij*_), neuromodulators).

##### Updated Synaptic Weight Equation

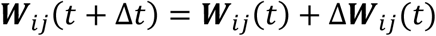

#### (3.3) Integrating into the Population Dynamics Model Full Model Equations

1. **Neural Activity Dynamics:**

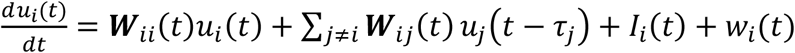
2. **Synaptic Plasticity Dynamics:**

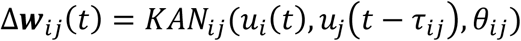
3. **Synaptic Weight Update**:

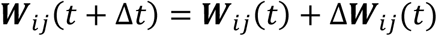

### (4.) Detailed Explanation and Implementation

#### (4.1) Selection of Variables for KANs

- Input Variables 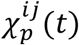:
  ∘ Presynaptic Activity *u*_*j*_(*t − τ*_*ij*_)
  ∘ ***Postsyna***ptic Activity *u*_*i*_(*t*)
  ∘ ***Neuromod***ulators *θ*_*ij*_(e.g., dopamine levels)
  ∘ Temporal Factors *t* (time dependence)

#### (4.2) Designing the KAN for Synaptic Plasticity

##### Step 1: Define Univariate Functions

- 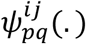:Functions applied to each input variable 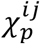.
- 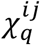: Output functions combining the summed inputs.

##### Step 2: Determine Network Architecture

- Layers:
  ∘ Input Layer: Receives 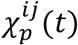.
  ∘ Hidden Layer: Applies 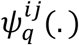 to each input.
  ∘ Summation Layer: Sums outputs of hidden layer.
  ∘ Output Layer: Applies 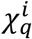 to the summed inputs and sums to get Δ*W*_*ij*_(*t*).

##### Step 3: Training the KAN

- **Objective:** Approximate the desired synaptic plasticity function *f*_*ij*_(*u*_*i*_(*t*), *u*_*j*_(*t − τ*_*j*_), *θ*_*ij*_).
- **Data Generation:** Use known plasticity rules (e.g., Hebbian, STDP) to generate training data.
- **Optimization:** Adjust parameters of 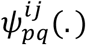 and 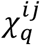 to minimize the error between the KAN output and 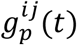.

#### (4.3) Implementation in the Multi-Area Model

- Time Integration:
  ∘ Discretize Time: Use a time step Δ*t*.
  ∘ At Each Time Step *t*:
    1. Compute Neural Activities *u*_*i*_(*t*).
    2. Compute Synaptic Changes Δ*w*_*ij*_(*t*) using KANs.
    3. Update Synaptic Weights *W*_*ij*_(*t* + Δ*t*).

Simulation Loop:

- For *t*= 0 to *T*:
  ∘ Update *u*_*i*_(*t*) and *W*_*ij*_(*t*)

### (5.) Implications and Applications

#### (5.1) Modeling Complex Plasticity Mechanisms

- **Flexibility:** KANs can model complex, nonlinear dependencies in synaptic plasticity that traditional rules might not capture.
- **Adaptability:** The network can learn from data, allowing it to model experimentally observed plasticity phenomena.

#### (5.2) Understanding Learning and Memory Formation

- **Multi-Area Interactions:** By incorporating KAN-modeled plasticity, we can simulate how learning in one brain area affects others.
- **Temporal Dynamics:** Capture how synaptic changes evolve over time with ongoing activity.

#### (5.3) Computational Efficiency

- **Dimensionality Reduction:** Simplifies the computation of synaptic updates in high-dimensional systems.
- **Scalability:** Suitable for large-scale simulations involving multiple brain regions.

### (6.) Example Scenario

**Objective:** Simulate learning across three interconnected brain areas with synaptic plasticity modeled by KANs.

#### (6.1) Model Setup

- **Areas:** A1 (primary), A2 (intermediate), A3 (higher-order).
- **Initial Conditions:** Random initial synaptic weights.
- **External Input:** Stimulus applied to A1.

#### (6.2) Simulation Steps

1. **Initialize Variables:**
  ∘ Set *u*_*i*_(0), KAN parameters.
2. **Time Loop:**
  ∘ For each *t*:
    ▪ **Compute t**.
    ▪ **Compute** Δ*W*_*ij*_(*t*) **using KANs**.
    ▪ **Update** *W*_*ij*_(*t* + Δ*t*).

#### (6.3) Observations

- **Synaptic Strengthening:** Observe how synaptic weights adapt based on activity patterns.
- **Inter-Area Coordination:** Analyze how changes in one area influence others.
- **Learning Outcomes:** Evaluate the network’s ability to learn and recognize patterns over time.

### (7.) Challenges and Considerations

#### (7.1) Determining KAN Functions

- **Function Selection:** Choosing appropriate univariate functions 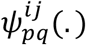 and 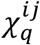 is critical.
- **Training Data:** Requires sufficient data representing the desired synaptic plasticity behaviors.

#### (7.2) Biological Plausibility

- **Interpretability:** KANs may be mathematically accurate but need to align with biological mechanisms.
- **Complexity vs. Simplicity:** Balancing model complexity with computational tractability and biological relevance.

#### (7.3) Computational Resources

- **Scalability:** Large networks with many areas may demand significant computational power.
- **Optimization:** Efficient algorithms are needed for training and simulation.

**Developing the Backpropagation Algorithm for Training the Kolmogorov-Arnold Network (KAN)**

**Prompt**

In the previous prompt, my notations were not clearly defined and were confusing. In this prompt, I further clear notations. Please develop the backpropagation algorithm for Training the KAN in 4.2 Step 3 again.

Assume that Nonlinear functions *f*_*p*_(*t*)_(*m,k*)_ for the *m*-th input variable and the *k*-th synapse define the interaction between input signals 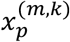 and synapses over time. To define the function *f* (*t*) for the *m*-th input variable as it propagates from an upstream brain region *k*_1_ to a downstream region *k*_*max*_, you sum or integrate these kernel functions across all synapses along the path.

The process of defining *f*_*p*_(*t*)_(*m*)_ involves integrating or summing up the effects of these nonlinear interactions as the input variable *m* propagates through the network, starting from the upstream brain region *k*_1_ and ending at the downstream brain region *k*_*max*_.

Mathematically, this could be represented as:

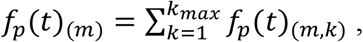

where *f*_*p*_(*t*)_(*m,k*)_ is the contribution of the *k*-th synapse to the overall function *f*_*p*_(*t*)_*m*_.

Define

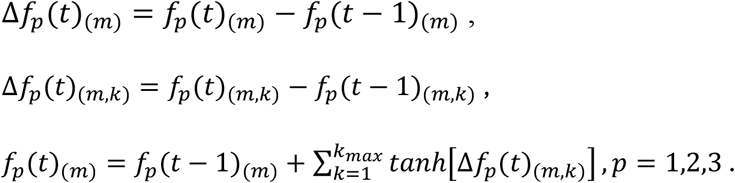

Let *h*(*t*)_(*m*)_ be actual output for the cumulative effect of the interaction between the *m*-th input variables and synapse along the paths from the upstream to downstream brain regions and *f*(*t*)_(*m*)_ be its corresponding desired output. Let 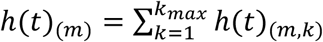.

***Define t***he energy function to be the square of the error between the desired output and actual output:

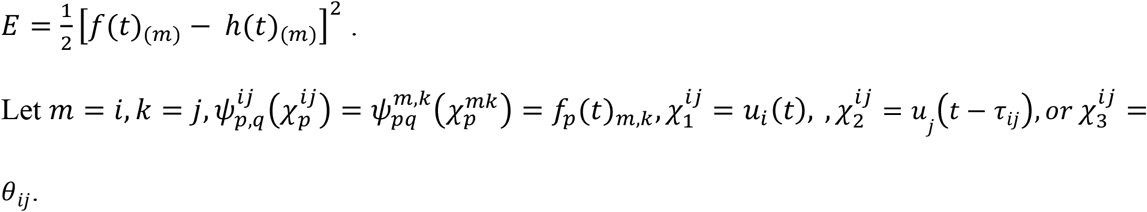

Assume that *f*_*ij*_(*u*_*i*_(*t*), *u*_*j*_(*t − τ*_*j*_), *θ*_*ij*_) in 3.2 synaptic plasticity models by KAN is replaced by

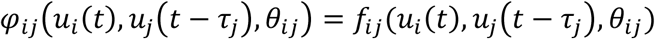

Please develop the backpropagation algorithm for Training the KAN in 4.2 Step 3.

**ChatGPTo1**

## BEGIN

**Thought for 62 seconds**

### Clarifying details

I’m piecing together the user’s request to redefine notations and enhance the backpropagation algorithm for training the KAN in 4.2 Step 3. This involves a thoughtful examination of the request’s specifics and drafting a compliant, helpful response.

### Revisiting definitions

Now I’m thinking through the latest definitions to ensure clarity in the backpropagation algorithm. This involves checking for compliant content and, if needed, providing accurate, step-by-step mathematical derivations.

### Clarifying notations

I’m mapping out the notations to ensure they are clear and consistent, which is crucial for understanding the backpropagation algorithm. This involves careful reading and precise definition.

### Revisiting backpropagation

I’m redoing the backpropagation algorithm for training the KAN, incorporating clearer notation definitions to enhance understanding. This involves refining how interactions between input signals and synapses are modeled, particularly for each input variable’s propagation through different brain regions.

### Charting the trajectory

I’m mapping out the function 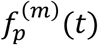 by integrating the nonlinear interactions from upstream to downstream brain regions, using sums and contributions from each synapse to describe the effects.

### Mapping out the backpropagation algorithm

I’m focusing on developing the backpropagation algorithm for training the KAN, using specified definitions including the energy function and synaptic plasticity models.

### Defining and updating

I’m outlining the steps to train the KAN by defining forward and backward passes, updating parameters based on error gradients, and minimizing the error between outputs.

### Clarifying the process

I’m ensuring variables are defined and consistent, then mapping out the forward pass, error computation, backward pass, and parameter updates, while avoiding policy violations.

### Mapping out the backpropagation

I’m detailing the backpropagation algorithm for the Kolmogorov-Arnold Network (KAN). The aim is to adjust KAN parameters, aligning actual output *h*^(*m*)^(*t*) with the desired output *f*^(*m*)^(*t*) by minimizing the energy function *E*.

### Mapping out cumulative functions

I’m detailing non-linear interactions, with the cumulative function 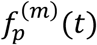 computed by summing individual synapse functions 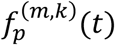.Adjustments and output analysis follow.

### Mapping forward pass

I’m detailing each step of the forward pass in a neural network, covering parameter computation, activation functions, and cumulative functions to achieve the final output *h*^(*m*)^(*t*).

### Calculating gradients

I’m mapping out gradients for updating parameters by computing error signals for activations, cumulative functions, and output weights. Each step, including computing gradients for synapses, is carefully considered to ensure precise adjustments.

### Refining weights

OK, let me see. I’m utilizing gradient descent to fine-tune the output and synaptic weights, ensuring each parameter adjustment steers the energy function minimization in the right direction.

### Understanding backpropagation

I’m digging into the forward and backward passes of the backpropagation algorithm. The forward pass involves calculating output and errors, while the backward pass computes gradients and updates parameters for synaptic and output weights.

### Navigating implementation

I’m making sure the learning rate, weight initialization, and temporal dependencies are spot on. Handling batch processing and activation function derivatives streamlines forward pass computations.

### Breaking down network dynamics

I’m detailing the forward pass, calculating changes in network states, and addressing how gradients are computed and applied during the backward pass for parameter updates. This evolution ensures synchrony in network development.

**END**

**Developing the Backpropagation Algorithm for Training the Kolmogorov-Arnold Network (KAN)**

## Introduction

In this section, we will develop the backpropagation algorithm for training the Kolmogorov-Arnold Network (KAN) based on your clarified notations. The goal is to adjust the parameters of the KAN so that the actual output *h*^(*m*)^(*t*) closely matches the desired output *f*^(*m*)^(*t*) by minimizing the energy function *E*.

### (1.) Definitions and Notations

#### (1.1) Nonlinear Interaction Functions

- 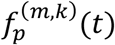: Nonlinear function for the *m*-th input variable and the *k*-th synapse at time *t*, defining the interaction between input signals 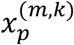 and synapses over time.
- **Cumulative Function** 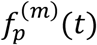: Sum of the nonlinear interactions across all synapses along the path from the upstream brain region *k*_1_ to the downstream brain region *k*_*max*_:

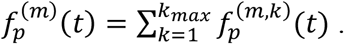

#### (1.2) Delta Functions

- **Change over Time for Cumulative Functions**:

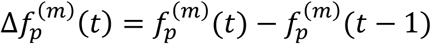
- **Change over Time for Individual Synapse Functions**:

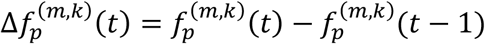

#### (1.3) Update Rule for Cumulative Function

- The cumulative function 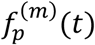 is updated as:

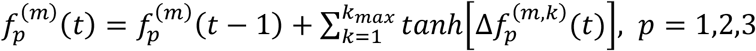

#### (1.4) Actual Output and Desired Output

- **Actual Output** *h*^(*m*)^(*t*): The cumulative effect of the interaction between *m*-th input variables and synapses along the paths from the upstream to downstream brain regions.
- **Desired Output** 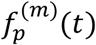: The target output we aim for.
- **Actual Output per Synapse**:

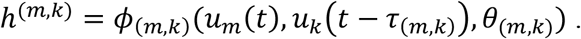

#### (1.5) Energy Function

- The energy function *E* is defined as:

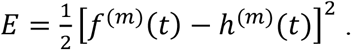

#### (1.6) Variable Substitutions

- **Indices**: *m*= *i, k*= *j*
- **Functions and Variables**:

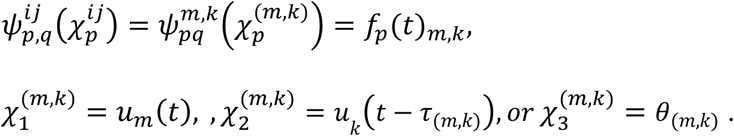
- **Synaptic Plasticity Functions**:

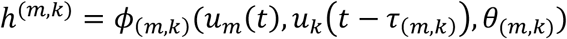

### (2.) Forward Pass Computation

The forward pass involves computing the outputs of the network given the inputs and current parameters.

#### (2.1) Compute 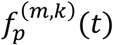

For simplicity, assume that 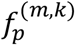 is a function of the input 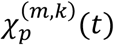 and parameters 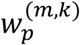:

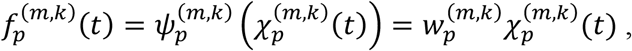

- 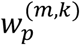: Weight parameter associated with the *p*-th input variable for the *m*-th input and *k*-th synapse.
- 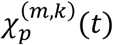 : Input variables, which are:

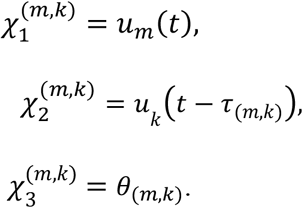

#### (2.2) Compute 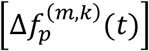

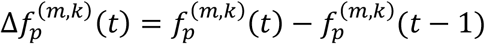

#### (2.3) Apply Activation Function

Compute the activation for each synapse:

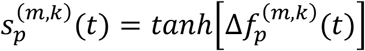

#### (2.4) Compute Cumulative Function 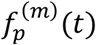

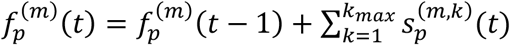

#### (2.5) Compute desired Output *h*^(*m*)^(*t*) and actual output *f*^(*m*)^(*t*)

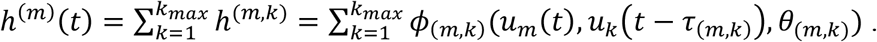

Assuming that the actual output *f*^(*m*)^(*t*) is a function of the cumulative functions 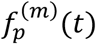:

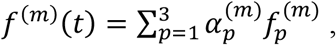

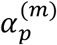: Output weight parameters combining the cumulative functions to produce the actual output.

#### (2.6) Compute Energy Function *E*

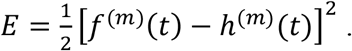

#### (3.1) Compute Error Signal at Output

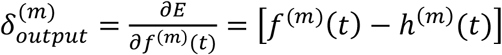

#### (3.2) Compute Gradients for Output Weights 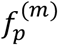

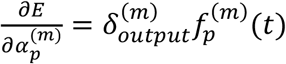

#### (3.3) Compute Error Signal for Cumulative Functions 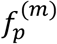

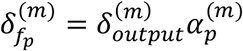

#### (3.4) Backpropagate Through the Summation

Since 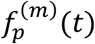 depends on 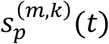:

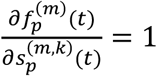

#### (3.5) Compute Error Signal for Activation Outputs 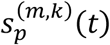

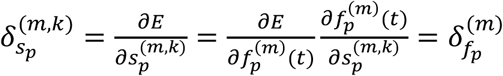

#### (3.6) Backpropagate Through Activation Function

Compute the derivative of the activation function:

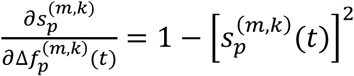

Compute the local gradient for each synapse:

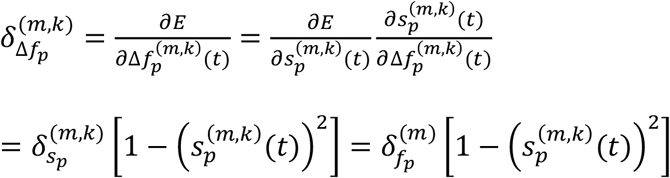

#### (3.7) Compute Gradient with Respect to 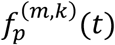

Since 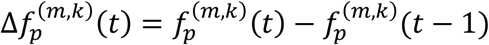

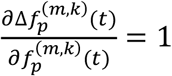

Therefore, the error signal for 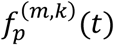 is

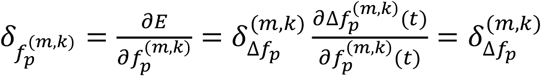

#### (3.8) Compute Gradient with Respect to Parameters 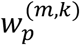

Since 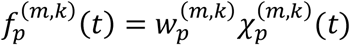,the gradient is

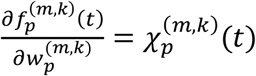

Thus, the gradient of the energy function with respect to 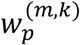 is

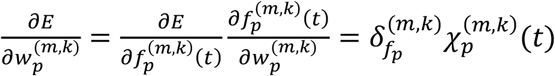

### (4.) Parameter Update Rule

Using gradient descent, update the parameters:

#### (4.1) Update Output Weights 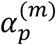

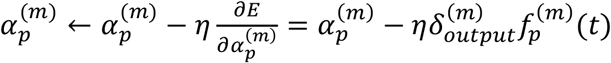

#### (4.2) Update Synaptic Weights 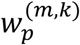

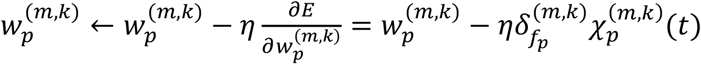

- *η*: Learning rate.

### (5.) Summary of the Backpropagation Algorithm

#### Forward Pass Steps

1. Compute Synaptic Functions:

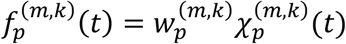
2. Compute Changes:

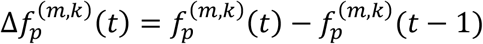
3. Apply Activation Function:

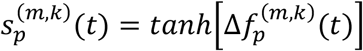
4. Compute Cumulative Functions:

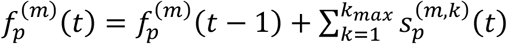
5. Compute Actual Output:

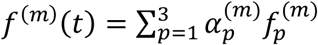
6. Compute Energy Function:

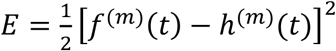

#### Backward Pass Steps

1. Compute Output Error Signal:

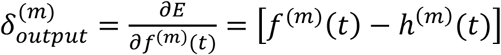
2. Compute Gradients for Output Weights:

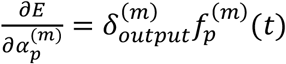
3. Compute Error Signals for Cumulative Functions:

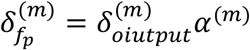
4. Compute Error Signals for Activation Outputs:

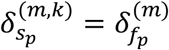
5. Compute Local Gradients:

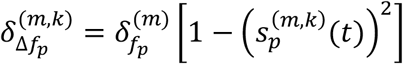
6. Compute Gradients for Synaptic Weights:

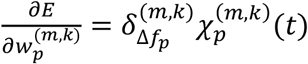
7. Update Parameters:
  - Output Weights:

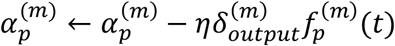

synaptic Weights:

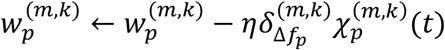

## (6.) Implementation Considerations

- **Learning Rate** *η*: Choose an appropriate learning rate to ensure convergence.
- **Initialization**: Initialize weights 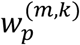 and 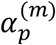 to small random values.
- **Temporal Dependencies**: Since 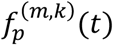 and 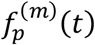 depend on their previous values, maintain a memory of past activations.
- **Batch Processing**: For improved performance, consider updating weights after processing a batch of training examples.
- **Activation Function Derivative**: Precompute *tanh* and its derivative to optimize computations.

### (7.) Example Calculation

Suppose:

- **Input Variables**:
  ∘ 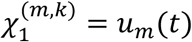
  ∘ 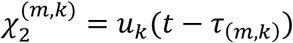
  ∘ 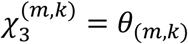
- **Initial Weights**: Initialize 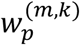 and 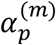 randomly.

#### Forward Pass

1. For each synapse *k* and input *p*:
  ∘ Compute 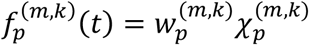.
2. Compute 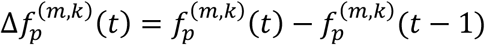.
3. Apply activation function:

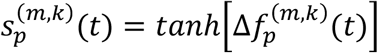
4. Sum over all synapses to get 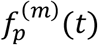.
5. Compute actual output:

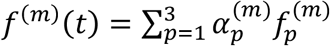
6. Compute the energy *E*.

#### Backward Pass

1. Compute output error signal 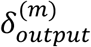.
2. Compute gradients for output weights 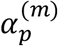.
3. Compute error signals 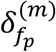.
4. For each synapse *k* and input *p*:
  ∘ Compute local gradients 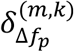.
  ∘ Compute gradients 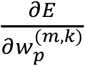
5. Update parameters 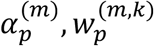.

Repeat these steps for each time step *t* and iterate until convergence.

### (8.) Summary

By developing the backpropagation algorithm as outlined, we can train the Kolmogorov-Arnold Network (KAN) to model synaptic plasticity within the multi-area neural population dynamics framework. This approach allows the network to adjust synaptic weights based on the error between the desired and actual outputs, enabling learning and adaptation over time.

**Mathematical Formula for Describing Dynamic Population Models Acting on Decoding Thought**

**Prompt**

**Please give a specific mathematical formula for describing dynamic population models acting on Decoding Thought**.

## ChatGPTo1

**Thought for 18 seconds**

**BEGIN**

### Considering policy compliance

I’m figuring out that I need to ensure the mathematical formula for dynamic population models is accurate, helpful, and aligns with OpenAI’s guidelines. Wouldn’t want errors or inappropriate content to creep in!

### Introducing neural models

I’m working through state-space or neural mass models to represent neural activity and decoding thought, focusing on dynamic neural field equations and recurrent neural networks.

### Formulating neural dynamics

I’m developing a model of neural population dynamics with differential equations for firing rates, including synaptic weights and external inputs.

### Understanding neuron connections

OK, let me see. Element *W*_*ij*_ reflects the link from population jjj to population iii, while *ϕ*(***r***(*t*)) applies a nonlinear function to firing rates.

### Connecting neural dynamics

I’m thinking through how neural dynamics are captured with differential equations, considering factors like firing rates, external inputs, and nonlinear activation. This leads to the development of the observation model.

### Decoding specific thoughts

I’m curious about how unique cognitive states link to patterns in ***r***(*t*). Using filtering methods to estimate these states and machine learning for classification or regression could work well.

### Decoding motor intentions

I’m piecing together how neural populations in the motor cortex can represent imagined hand movements. Using data like sensory cues and neural signals, I estimate thought states and train classifiers for decoding neural signals to intentions.

### Enhancing real-time decoding

I’m digging into adding synaptic plasticity and manifold representation to ameliorate decoding of motor intentions, using variable learning rates and manifold learning techniques to capture the low-dimensional structure within high-dimensional data.

**END**

## Introduction

Decoding thought involves interpreting neural activity to infer underlying cognitive processes. Dynamic population models provide a mathematical framework to describe how neural populations evolve over time and how their collective dynamics correspond to specific thoughts or intentions. One such model is the **neural population firing rate model**, which captures the temporal evolution of neural firing rates in response to synaptic inputs and external stimuli.

### Mathematical Model

The dynamics of the neural population can be described by the following differential equations:

1. **Neural Dynamics Equation**

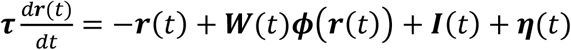
2. **Observation Equation**

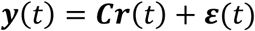

### Explanation of Variables and Parameters

1. **Neural Dynamics Equation Variables:**
  ∘ ***r***(*t*): An NNN-dimensional vector representing the firing rates of NNN neural populations at time *t*. Each element *r*_*i*_(*t*) corresponds to the firing rate of neural population *i*.
  ∘ *τ*: A time constant representing the characteristic response time of the neural populations.
  ∘ ***W***: An *N*× *N*synaptic weight matrix. The element *W*_*ij*_ represents the strength and influence of neural population *j* on population *i*.
  ∘ ∅(***r***(*t*)): A nonlinear activation function applied element-wise to ***r***(*t*). Common choices include:
    ▪ **Sigmoid function**

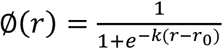
    ▪ **Hyperbolic tangent**:

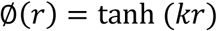
    ▪ **Rectified Linear Unit (ReLU)**:

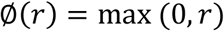
    ▪ **In the vector form**:

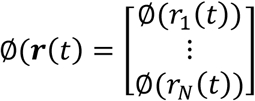
  ∘ ***I***(*t*): An *N*-dimensional external input vector representing stimuli or inputs affecting the neural populations.
  ∘ ***η***(*t*): An *N*-dimensional process noise vector accounting for random fluctuations and unmodeled dynamics.
2. **Observation Equation Variables:**
  ∘ ***y***(*t*): An *M*-dimensional vector of observed signals at time *t*, such as measurements from EEG, MEG, or intracortical recordings.
  ∘ ***C***: An *M* × *N* observation matrix mapping the neural firing rates to the observed signals. The element *C*_*ij*_represents the contribution of neural population *j* to observation *i*.
  ∘ ***ε***(*t*): An *M*-dimensional observation noise vector representing measurement noise and errors.

### Decoding Thought Using the Model

1. **State Representation**:
  ∘ The firing rates *r*(*t*) represent the internal neural state associated with specific thoughts or cognitive processes.
2. **Dynamics Modeling**:
  ∘ The neural dynamics equation models how these firing rates evolve over time due to intrinsic neural interactions (***W***∅(***r***(*t*))) and external inputs (***I***(*t*)).
3. **Observation Mapping**:
  - The observation equation relates the internal neural states ***r***(*t*) to measurable signals ***y***(*t*).
4. **Decoding Process**:
  - **State Estimation**: Use filtering techniques (e.g., Extended Kalman Filter, Particle Filter) to estimate ***r***(*t*) from ***y***(*t*).
  - **Classification/Regression**: Apply machine learning algorithms to the estimated ***r***(*t*) to decode the thought content or predict cognitive variables.

### Incorporating Synaptic Plasticity

To model learning and adaptation, synaptic plasticity can be integrated into the synaptic weight matrix ***W***(*t*):

### Synaptic Plasticity Equation

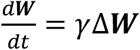

- *γ*: Learning rate controlling the speed of synaptic changes.
- Δ***W*** : Synaptic modification term, which can be modeled using Hebbian learning rules or more complex functions like those provided by Kolmogorov-Arnold Networks (KANs).

### Kolmogorov-Arnold Networks for Synaptic Plasticity

- **Purpose**: Approximate complex, multivariate synaptic update functions.
- **Synaptic Modification Term**:

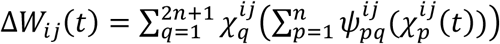
- 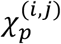: Inputs influencing synaptic changes, such as pre- and postsynaptic firing rates *r*_*j*_(*t*), *r*_*i*_(*t*), time delays, or neuromodulators.
- *ψ*_*pq*_, *χ*_*q*_: Univariate functions forming the KAN architecture.

### Full Model Including Synaptic Plasticity

1. **Neural Dynamics with Time-Varying Weights**

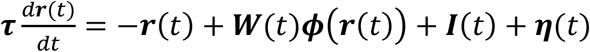
2. **Synaptic Weight Updates**

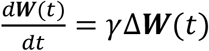
  - With Δ***W***(*t*)computed usin a KANs.
3. **Observation Equation**

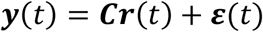

### Decoding Algorithm Steps

1. **Data Collection**:
  ∘ Record observations ***y***(*t*) while subjects engage in specific thought processes.
2. **Initialization**:
  ∘ Initialize model parameters ***W***(0), ***C***, and neural states ***r***(0)
3. **State Estimation**:
  ∘ Use filtering techniques to estimate ***r***(*t*) from ***y***(*t*).
4. **Synaptic Update**:
  ∘ Update ***W***(*t*) using the synaptic plasticity equation and KANs.
5. **Decoding Thought**:
  ∘ Apply machine learning models (e.g., classifiers) to ***r***(*t*) to predict the thought content.
6. **Learning and Adaptation**:
  ∘ Continuously update the model parameters to improve decoding accuracy over time.

### Example Application: Decoding Visual Imagery

Suppose we aim to decode whether a subject is imagining a specific visual scene:

1. **Neural Populations**:
  ∘ Represent neural groups in the visual cortex.
2. **External Input**:
  ∘ Visual cues or instructions prompting the subject to imagine certain scenes.
3. **Data Collection**:
  ∘ Record EEG or fMRI signals ***y***(*t*) during the task.
4. **Model Implementation**:
  ∘ Use the dynamic model to estimate ***r***(*t*).
5. **Synaptic Plasticity**:
  ∘ Model how synaptic connections ***W***(*t*) adapt during the imagination process.
6. **Decoding**:
  ∘ Classify ***r***(*t*) to determine the content of the visual imagery.

### Mathematical Considerations

- **Nonlinear Dynamics**:
  ∘ The activation function ∅ introduces nonlinearity, essential for capturing complex neural behavior.
- **Stochastic Elements**:
  ∘ Process noise ***η***(*t*) and observation noise ***ε***(*t*) account for variability and uncertainties.
- **Parameter Estimation**:
  ∘ Parameters (*t*), ***C***, and those within KANs are learned from data using optimization techniques like gradient descent or expectation-maximization.

### Benefits of the Model

- **Captures Temporal Dynamics**:
  ∘ Models the evolution of neural activity over time, essential for decoding thoughts that are dynamic in nature.
- **Incorporates Learning**:
  ∘ Synaptic plasticity allows the model to adapt, mirroring learning processes in the brain.
- **Handles Nonlinearity**:
  ∘ Nonlinear activation functions and KANs enable the model to capture complex relationships.

### Challenges and Considerations

- **Model Complexity**:
  ∘ Requires careful tuning of parameters and may be computationally intensive.
- **Data Quality**:
  ∘ Accurate decoding depends on high-quality neural recordings and sufficient data for training.
- **Individual Variability**:
  ∘ Neural dynamics can vary significantly between individuals; personalized models may be necessary.

### Summary

The provided mathematical model offers a framework for describing dynamic population models acting on decoding thought. Its diagram is shown in Figure 9. By modeling neural population dynamics, incorporating synaptic plasticity, and relating internal neural states to observable signals, we can develop algorithms capable of inferring thought processes from neural data. This approach has significant implications for neuroscience research and practical applications like brain-computer interfaces.

## Discussion

To demonstrate the ability of ChatGPTo1 for scientific discovery, we introduced our first model: the “Dynamic Models of Neural Population Dynamics.” ChatGPTo1, after describing the widely used linear state-space models of neural population dynamics, including observation models, combined with ChatGPT4o to implement numerical simulations of these linear state-space models. Subsequently, I used prompts to require ChatGPTo1 to develop a multi-area nonlinear neural dynamical model from upstream brain regions to downstream brain regions. Then, through prompts, I asked to incorporate synaptic plasticity mechanisms used to simulate learning and adaptation into the above model. Finally, I requested that ChatGPTo1 integrate synaptic plasticity modeled by Kolmogorov-Arnold networks into multi-area neural population dynamics models. Incorporating this nonlinear relationship of synaptic plasticity modeled by Kolmogorov-Arnold networks into multi-area nonlinear neural population dynamics models is entirely new, not found in the literature, and impossible to establish through pre-training methods. Furthermore, we asked ChatGPTo1 to further include the backpropagation (BP) algorithm used to train the Kolmogorov-Arnold networks into the model. However, because providing the specific model structure of the Kolmogorov-Arnold networks is very complex, the mathematical symbols in the first prompt were inconsistent. In reviewing the results, we found that some parts of the nonlinear dynamical model had errors. After we corrected the errors due to symbol inconsistencies, except for one symbol error, the rest of the system gave correct results. The biggest deficiency of ChatGPTo1’s model is that it cannot provide an end-to-end reasoning model, cannot fully automate multi-stage logical reasoning, and cannot implement data analysis and simulation calculations, thus lacking the ability to check whether the reasoning is correct. Currently, ChatGPTo1 cannot process image, audio, and video multimodal data, nor can it effectively perform complex data analysis. However, ChatGPT4o has strong capabilities in handling multimodal data, including images. The system provides functions for mutual switching, which to some extent extends ChatGPTo1’s capabilities, but the switching is not very smooth. The core of scientific research is innovation: to produce new concepts, hypotheses, and coherent theories based on original theories and new observational data. Innovation mainly relies on reasoning. Logical reasoning includes serial reasoning like “chain-of-thought,” tree-like chain-of-thought, mind maps, thinking frameworks, procedural thinking, and so on. These new thinking patterns should also be included in future models.

## Notes

### Competing Interest Statement

The authors have declared no competing interest.

## References

1. Si, C., Yang, D., Hashimoto, T. (2024). Can LLMs Generate Novel Research Ideas? A Large-Scale Human Study with 100+ NLP Researchers. 2409.04109.

2. Lee, K., et al. (2020). BioBERT: a pre-trained biomedical language representation model for biomedical text mining. Bioinformatics, 36(4), 1234–1240.

3. Wang, L. L., et al. (2019). CORD-19: The COVID-19 Open Research Dataset. ArXiv preprint, 2004.10706.

4. Surden, H. (2019). Artificial intelligence and law: An overview. Georgia State University Law Review, 35(4), 1305.

5. Cohen, P. R. (2015). DARPA’s Big Mechanism program. Physical Biology, 12(4), 045008.

6. Jumper, J., et al. (2021). Highly accurate protein structure prediction with AlphaFold. Nature, 596(7873), 583–589.

7. Esteva, A., et al. (2017). Dermatologist-level classification of skin cancer with deep neural networks. Nature, 542(7639), 115–118.

8. Delépine, M., et al. (2022). Domain-specific language model pretraining for biomedical natural language processing. Journal of Biomedical Informatics, 128, 104052.

9. Wang, P., et al. (2021). Augmented intelligence: The next frontier in AI. IEEE Transactions on Artificial Intelligence, 2(6), 445–452.

10. Delépine, M., et al. (2022). Domain-specific language model pretraining for biomedical natural language processing. Journal of Biomedical Informatics, 128, 104052.

11. Krishna V. Shenoy, E.V., and Kao, J. C. (2021). Measurement, manipulation and modeling of brain-wide neural population dynamics. Nat Commun 12, 633 (2021).

12. Bittner, S. R. et al. (2017). Population activity structure of excitatory and inhibitory neurons. PLoS One. 2017; 12(8): e0181773.

13. Friston, K. Dynamic Causal Modeling for EEG and MEG.

14. Box, G. E. P. et al. Time Series Analysis: Forecasting and Control.

15. Friston, K.J., Harrison, L., & Penny, W. (2003). Dynamic causal modelling. NeuroImage, 19(4), 1273–1302.

16. Baillet, S., Mosher, J.C., & Leahy, R.M. (2001). Electromagnetic brain.

17. Appelbaum, L. G., Shenasa, M. A., Louise Stolz, L., Daskalakis, Z. (2023). Synaptic plasticity and mental health: methods, challenges and opportunities. Neuropsychopharmacol. 48, 113–120 (2023).

18. Weishaupt, N. (2017). Chapter 2: Cortical Plasticity in Response to Injury and Disease, in The Cerebral Cortex in Neurodegenerative and Neuropsychiatric Disorders.

19. Tong, R., Nigel J. Emptage, N.J., Padamsey, Z. (2020). A two-compartment model of synaptic computation and plasticity. Mol Brain 13, 79 (2020).

20. Andrade-Talavera, Y., Fisahn, A. & Rodríguez-Moreno, A. Timing to be precise? An overview of spike timing-dependent plasticity, brain rhythmicity, and glial cells interplay within neuronal circuits. Mol Psychiatry 28, 2177–2188 (2023).

21. Kavalali, E.T., Monteggia, L.M. Rapid homeostatic plasticity and neuropsychiatric therapeutics. Neuropsychopharmacol. 48, 54–60 (2023).

22. Theoretical Neuroscience” by Peter Dayan and L.F. Abbott.

23. “Neuronal Dynamics” by Wulfram Gerstner, Werner M. Kistler, Richard Naud, and Liam Paninski.

24. Markram, H., Lübke, J., Frotscher, M., & Sakmann, B. (1997). Regulation of synaptic efficacy by coincidence of postsynaptic APs and EPSPs. Science, 275(5297), 213–215.

25. Morrison, A., Aertsen, A., & Diesmann, M. (2007). Spike-timing-dependent plasticity in balanced random networks. Neural Computation, 19(6), 1437–1467.

26. Lillicrap, T.P., Santoro, A., Marris, L., Akerman, C.J., & Hinton, G. (2020). Backpropagation and the brain. Nature Reviews Neuroscience, 21(6), 335–346.

27. Richards, B.A., Lillicrap, T.P., et al. (2019). A deep learning framework for neuroscience. Nature Neuroscience, 22(11), 1761–1770.

28. Kavalali ET, Monteggia LM. Targeting Homeostatic Synaptic Plasticity for Treatment of Mood Disorders. Neuron. 2020 Jun 3;106(5):715–726.

